# Dysregulation of mTOR Signaling Mediates Common Neurite and Migration Defects in Both Idiopathic and 16p11.2 Deletion Autism Neural Precursor Cells

**DOI:** 10.1101/2022.09.17.508382

**Authors:** Smrithi Prem, Bharati Dev, Cynthia Peng, Monal Mehta, Rohan Alibutud, Robert J. Connacher, Madeline St Thomas, Xiaofeng Zhou, Paul Matteson, Jinchuan Xing, James H. Millonig, Emanuel DiCicco-Bloom

## Abstract

Autism spectrum disorder (ASD) is defined by common behavioral characteristics, raising the possibility of shared pathogenic mechanisms. Yet, vast clinical and etiological heterogeneity suggests personalized phenotypes. Surprisingly, our iPSC studies find that six individuals from two distinct ASD-subtypes, idiopathic and 16p11.2 deletion, have common reductions in neural precursor cell (NPC) neurite outgrowth and migration even though whole genome sequencing demonstrates no genetic overlap between the datasets. To identify signaling differences that may contribute to these developmental defects, an unbiased phospho-(p)-proteome screen was performed. Surprisingly despite the genetic heterogeneity, hundreds of shared p-peptides were identified between autism subtypes including the mTOR pathway. mTOR signaling alterations were confirmed in all NPCs across both ASD-subtypes, and mTOR modulation rescued ASD phenotypes and reproduced autism defects in controls. Thus, our studies demonstrate that genetically distinct ASD subtypes have common defects in neurite outgrowth and migration which are driven by the shared pathogenic mechanism of mTOR signaling dysregulation.

## INTRODUCTION

Autism spectrum disorders (ASD) are characterized by deficits in social interaction and communication and the presence of repetitive and restrictive interests and behaviors. Though these common impairments define ASD, there is marked heterogeneity in both clinical presentation and etiological underpinnings amongst affected individuals (Jones W. et al., 2009; Kim S. H. et al., 2013; Betancur C., 2011). In turn, this heterogeneity raises the possibility that there are “different” ASD subtypes (Ousley O. et al., 2014; Eaves L. C. et al., 1994; Beglinger L. J. et al., 2001). Yet, despite this heterogeneity, all autism subtypes are united by their core behavioral phenotypes. Consequently, we postulate that there may be common pathways linking different autism subtypes. Yet, few studies have compared multiple ASD subtypes to investigate possible commonalities.

About 10-15% of autism cases are genetically-defined and caused by single gene mutations or copy number variants (CNVs) while most of ASD etiology remains undefined or idiopathic. Yet, much of our knowledge of ASD pathophysiology is derived from rodent studies of syndromic rare variant genes. Consequently, our insight into a majority of ASD pathophysiology has been limited and we have little knowledge of the similarities and differences between idiopathic and genetically-defined autisms. Several studies have uncovered the convergence of both rare and common variant ASD risk genes onto the developing mid-fetal cerebral cortex (8-24 week) (Willsey A. J. et al., 2013; Parikshak N. N. et al., 2013; Satterstrom F. K. et al., 2020; Grove J. et al., 2019). During this period, neural precursor cells (NPCs) undergo proliferation, migration, and differentiation to form neurons and establish normal brain cytoarchitecture (Liszewska E. et al., 2018). Now, with the advent of induced pluripotent stem cell (iPSC) technology, we can study human neural development in cells derived from individuals with both genetic and idiopathic neuropsychiatric disorders. Thus far, iPSC studies of monogenic, CNV, and idiopathic ASD have observed alterations in basic developmental processes, such as synapse formation (Acab A. et al., 2015; Yang G. et al., 2020). However, many of these studies focused on terminally differentiated neurons, with very few examining developmental events that correspond to mid-fetal cortical development that precede synapse formation, such as neurite outgrowth and cell migration (Packer A., 2016). Moreover, there is little insight into whether and how extracellular factors (EFs), which coordinate neurodevelopment by regulating signaling, contribute to ASD pathogenesis in human NPCs.

To begin addressing these questions, we studied early neurite development and cell migration using human iPSC derived-NPCs from two different autism subtypes–an idiopathic cohort (I-ASD) and a genetically defined CNV (16p11.2 deletion) cohort. We demonstrate remarkable convergence in neurite and migration phenotypes despite whole genome sequencing (WGS) showing no genetic overlap between the datasets. To identify mediating signaling pathways, an unbiased p-proteome was conducted, and it defined a significant overlap in p-peptides between the datasets. Bioinformatics revealed the mTOR pathway as a point of convergence between the two subtypes, which was supported by western studies. Further, modulation of the mTOR pathway with small molecules rescued and reproduced ASD phenotypes indicating that mTOR signaling defects are mechanistically linked to the neurodevelopmental phenotypes. Nonetheless, there were also ASD subtypes as evidenced by differential responses to EFs as well as distinct mTOR alterations including elevations and reductions. Overall, our studies indicate that despite the genetic and phenotypic heterogeneity two distinct ASD subtypes have common defects in mTOR signaling which mediate the surprisingly common neurodevelopmental phenotypes.

## MATERIALS AND METHODS

### Patient Cohorts

#### Idiopathic Autism (I-ASD)

I-ASD patients were selected from a larger cohort of 85 New Jersey families recruited by the Brzustowicz laboratory as part of the New Jersey Language and Autism Genetics Study (NJLAGS). In these families, at least one individual meets the criteria for autism spectrum disorder (ASD) and at least one other family member meets criteria for language-based learning impairment (LLI). Each family member was extensively phenotyped by the same set of clinicians with a battery of behavioral tests as described in Bartlett et al 2012 and 2014. The ASD proband was required to meet the criteria for “Autistic Disorder” on two of the three following measures: 1) Autism Diagnostic Interview-Revised (ADI-R), 2) Autism Diagnostic Observation Scale (ADOS), and Diagnostic and Statistical Manal IV (DSM-IV). Moreover, the ASD individual had no known genetic causes of autism such as Fragile X or Rett Syndrome. The LLI individual was identified by ruling out ASD, hearing impairments, and other neurological disorders and by using multiple language assessments including the Comprehensive Test of Language Fundamentals (Bartlett C. W. et al., 2012; Bartlett C. W. et al., 2014). In addition, each family member underwent 3 to 5 hours of direct behavioral testing including members of the family who do not have LLI or ASD. All family members were also assessed with the DSM-IV and ADOS to ensure that unaffected siblings and LLI did not meet criteria for autism. From this broader cohort, we selected 8 families with the following characteristics: 1) ASD proband with moderate or severe symptoms 2) families with sex matched unaffected Sib (without ASD or LLI). The studies presented are on 3 families from this smaller cohort of 8. Blood samples were obtained from all members of these families and lymphocytes were cryopreserved were used to generate iPSCs.

For clinical characterization of our I-ASD cohort, please refer to the latest publication from our lab which features clinical data for each individual (Connacher et al 2022, Figure 1). Briefly, I-ASD-1 has severe cognitive impairment (unable to complete IQ test) and ADOS and ADI-R revealed limited language comprehension to a few single words as well as occasional echolalic and scripted speech. SRS score was 90 indicating severe social impairment. I-ASD-1 had 78^th^ percentile head circumference at 4.1 years indicating normal head size. I-ASD-2 also has severe cognitive impairment (unable to complete IQ test) with ADOS and ADI-R revealing limited language comprehension to single words and directions and almost no language production. SRS score was 83 (severe social impairment) and proband had a head circumference of 97^th^ percentile at time of measurement (14 years) consistent with macrocephaly. Lastly, I-ASD-3 has limited language comprehension to a small number of single words, Non-verbal IQ (NVIQ) of 118 and SRS score of 69 (moderate social impairment). Unaffected individuals were diagnostically determined to not have any language or learning impairment or ASD as noted above.

**Figure 1:**
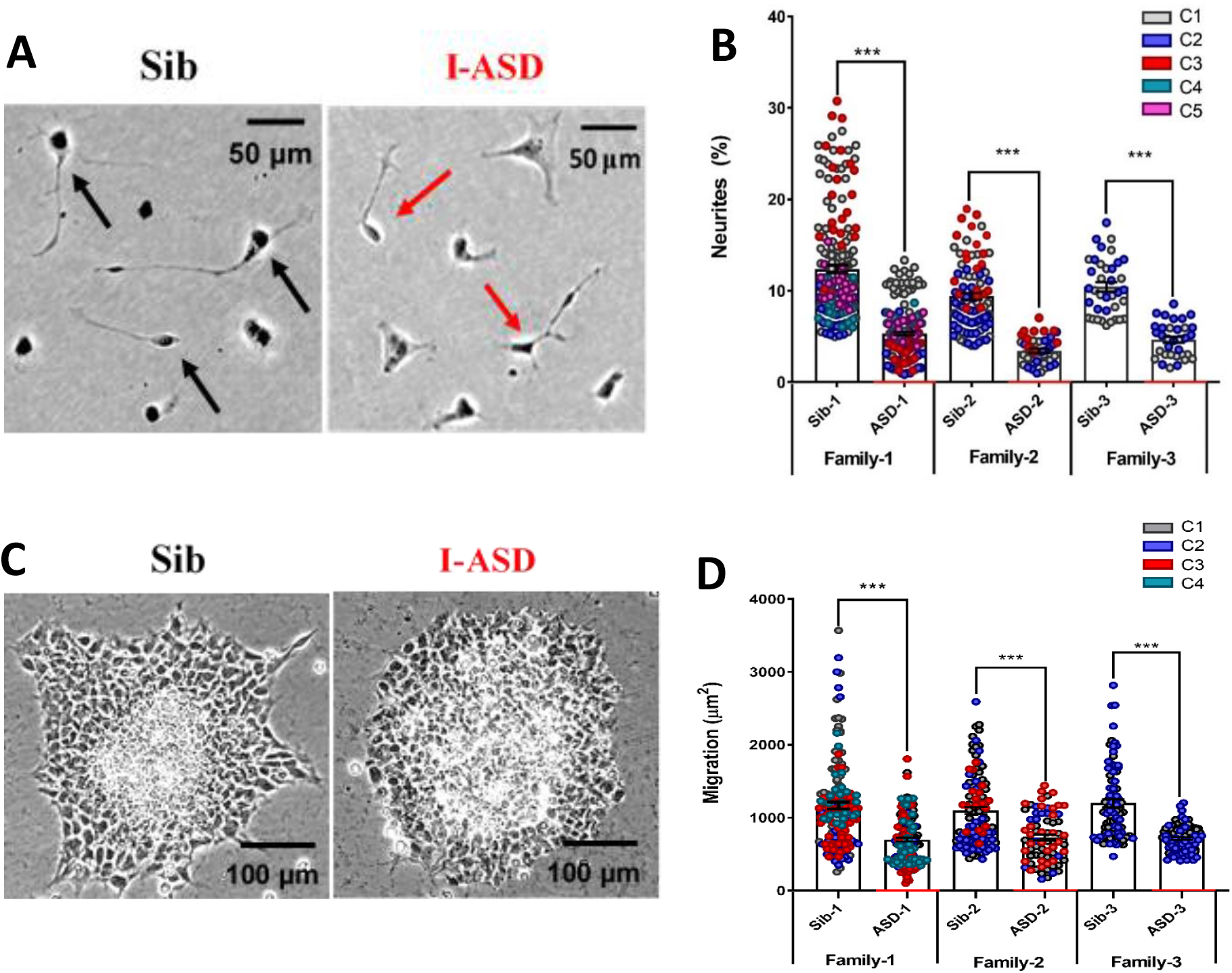
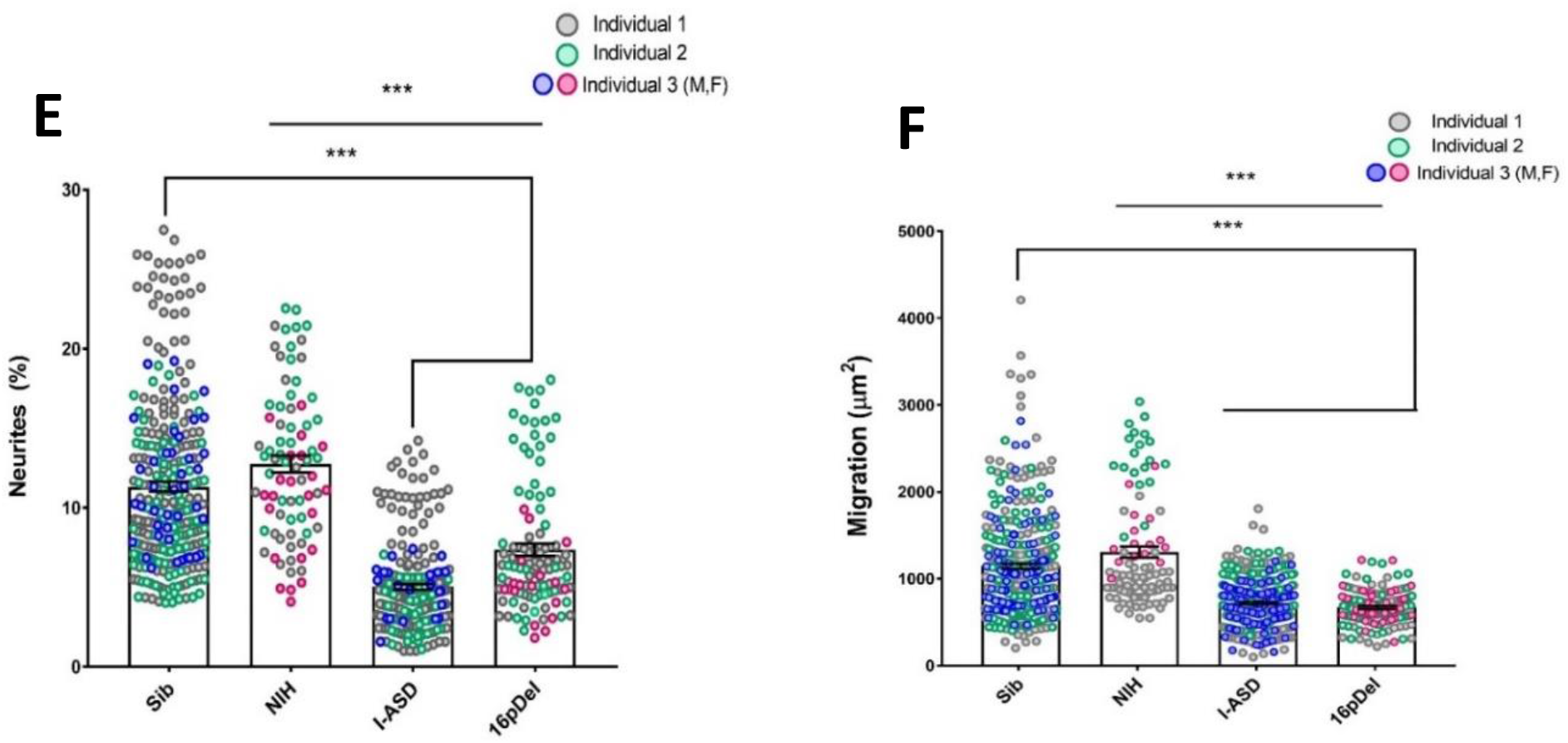
Reduced Neurite outgrowth and Cell Migration in both I-ASD and 16pDel NPCs. A) Representative image: neurite outgrowth at 48h showing more cells with neurites in Sib (black arrows) than I-ASD (red arrows). B) Quantification of % neurites in 3 pairs of I-ASD and Sibs: I-ASD NPCs have fewer neurites (%) than Sib NPCs in all families. In scatterplot graphs of neurite outgrowth, each dot represents the % of cells with neurites in a single dish. Colors represent different clones. C) Representative image showing Sib neurosphere with migrating carpet of cells (dark) moving further than that of the I-ASD neurosphere D) Cell migration in 3 pairs of I-ASD and Sibs: I-ASD NPCs migrate less than Sib NPCs in all families. In scatter plot graphs of cell migration, each dot represents migration of an individual neurosphere. All sib vs I-ASD comparisons: Student’s T-test E and F) Neurite outgrowth (E) and migration (F) in Sibs, NIH controls, I-ASD and 16pDel ASD. Each dot represents the data from a single dish with different colors denoting different individuals. Both 16pDel and I-ASD have reduced neurite outgrowth and migration compared to Sib and NIH NPCs. F) NPC migration in Sibs, NIH, I-ASD, and 16pDel ASD. Both 16pDel and I-ASD NPCs have reduced migration compared to both Sib and NIH. 1-way ANOVA for all 16pDel experiments with P<0.001 for all unaffected vs affected comparisons.

#### 16p11.2 deletion and NIH controls

Fibroblasts and lymphocytes were acquired from two males and one female patient with the 16p11.2 deletion and autism. These individuals were derived from the Simon’s Foundation VIP cohort, now termed Simon Searchlight Collection. Identity codes from SFARI for the individuals are as follows Female (14758.x3), Male-1 (14824.x13), and Male-2 (14799.x1). ASD inclusion criteria for 16pDel probands required they meet ADI-R and ADOS score cutoff criterion for autism spectrum disorder or autism (some individuals were clinically assessed using DSM-IV criteria). The probands were assessed on verbal and nonverbal cognitive abilities as described previously (Simons VIP, 2012). As we reported in Connacher et al., 2022, the first male (16pDel-1) exhibited autism spectrum disorder (Asperger’s Disorder), with a full-scale IQ (FSIQ) of 122, (non-verbal IQ [NVIQ] 130, verbal IQ [VIQ] 106), head circumference at the 99^th^ percentile at 14.5 years, consistent with macrocephaly, and comorbid expressive language disorder, anxiety, and microphthalmia with an SRS score of 76. The second male (16pDel-2) exhibited autism, with FSIQ score of 93 (NVIQ 98, VIQ 87) and a head circumference at the 99^th^ percentile at age 14.3 years, consistent with macrocephaly. He also displayed numerous other developmental phenotypes including coordination disability, developmental delay, cerebral palsy, ADD/ADHD, articulation disorder, and repetitive /expressive language disorder and a SRS score of 90. Lastly 16pDel-F exhibited atypical autism (previously PDD-NOD) with a full scale IQ (FSIQ) of 71 and NVIQ of 82, with a head circumference of 52.5 at 7.2 years (80^th^ percentile) as well as comorbid articulation disorder, communication disorder, and ADHD.

For 16pDel two iPSC clones per individual were obtained from RUCDR Infinite Biologics (now Infinity Biologix). As genetically matched siblings were unavailable for 16pDel, three sex matched iPSC control lines from genetically normal newborns from the NIH Regenerative Medicine Program were obtained. Specifically, 2 male (NCRM-1 & NCRM-3) and 1 female (NCRM-6) research grade line were utilized. The NIH control iPSC were generated from cord blood using episomal plasmid reprogramming method. https://commonfund.nih.gov/stemcells/lines. 16pDel were also compared to the Sibs from I-ASD cohort.

### Derivation, Validation, and Maintenance of iPSC lines

Sib and I-ASD iPSCs were generated by the Lu and Millonig labs while the 16pdel iPSCs were generated by RUCDR. For I-ASD and Sib, T lymphocytes were isolated from cryopreserved samples and these cells were expanded and then reprogrammed with a non-integrating Sendai virus containing the 4 Yamanaka factors: SOX2, OCT3/4, KLF, and a temperature sensitive C-MYC (Seki T. et al., 2011). The 16pdel iPSCs were also made using the same Sendai virus method however, both lymphocytes and fibroblasts were used as the initial somatic tissue. iPSCs were characterized using immunocytochemistry and QRTPCR for the following markers: NANOG, OCT4, TRA-1-60, SSEA4, CD24, and E-Cadherin. iPSCs were assayed for chromosomal abnormalities via G-band karyotype assay or via Array Comparative Genomic Hybridization (Company: Cell Line Genetics); DNA fingerprinting was also conducted to confirm genetic identity to original tissue samples. See Connacher et al 2022 (in press, Stem Cell Reports), materials and methods and supplementary information section for details and data on these iPSC characteristics. iPSC lines were maintained in mTeSR™1 media (Stem Cell Technologies, 85850) on 6-well plates (Corning, COR-3506) coated with hESC qualified Matrigel (Corning, 354277). Media was changed daily. Once cells reached 70-90% confluency, they were passaged by incubating cells with 0.5 mM filter-sterilized EDTA diluted in 1X PBS for 10-20 mins. When iPSCs detached from the plate, they were lifted and centrifuged at 150g for 5 mins. The cell pellet was then gently resuspended in mTeSR and plated at 15-30% confluence in mTeSR media with 5uM rock inhibitor (Y-compound), Y27632 (Stem Cell Technologies, 72302) for 24 hours.

### Generation, Validation, and Maintenance of Neural Precursor Cells (NPCs)

To generate NPCs, iPSCs were induced using a modified version of the ThermoFisher GIBCO Neural Induction Protocol (Williams M. et al., 2018). In brief, iPSCs at 70-90% confluency were dissociated using 1X Accutase (ThermoFisher Scientific A111050) and were centrifuged at 150g for 5 minutes. The resulting pellet was then resuspended into Neural Induction Media (NIM) (ThermoFisher Scientific, A1647801) with 5uM of Y-compound. The following densities of iPSCs were plated into Matrigel coated 12-well plates containing 1mL of NIM (with 5μM Y-compound): 80K, 120K, 200K, 300K. Two wells of each density were made. One set of wells at each density were induced for 7 days while the other was induced for 8 days. Once induction day 7 or 8 was reached, cells were lifted with 1X Accutase and centrifuged at 300g and then resuspended in Neural Expansion Media (NEM) (ThermoFisher Scientific, A1647801). 1-1.5 million cells were then replated onto Matrigel-coated 6-well plate with 2 mL NEM with 5uM Y-compound. At this point cells are P0. For our studies, NPCs were used only from P3 to P8.

To assess NPC identity, immunocytochemistry (ICC) was conducted to verify that cells expressed the following NPC markers (Sox2, Pax6, and Nestin) NPCs were discarded if marker immunostaining revealed Nestin or Sox2 <85% or Pax6 < 60% (Connacher et al., 2022). NPCs were also stained with OCT3/4 to ensure they are no longer expressed this iPSC marker. As shown in Connacher et al 2022, Quantiplex analysis was also performed on NPC mRNA to assess expression of several NPC markers.

NPCs were also differentiated into neurons, astrocytes, and oligodendrocytes as per Thermofisher Protocol:https://www.thermofisher.com/us/en/home/references/protocols/neurobiology/neurobiology-protocols/differentiating-neural-stem-cells-into-neurons-and-glial-cells.html. Differentiated cell identity was assessed by ICC for the following markers: MAP2/Tau (Neuron), O4/Gal-c (oligodendrocytes) and GFAP for astrocytes (Connacher et al 2022).

### Whole Genome Sequencing and Variant Calling

Whole genome sequencing data for the samples in the I-ASD dataset was extracted from a previous study (Zhou et al, 2022). The details of sequencing and variant identification are described previously (Zhou et al, 2022). The sequencing data and the variant calls are available in the National Institute of Mental Health Data Archive (NDA) under projects C1932 and C2933.

The variants were annotated with ANNOVAR (ver. 2017-07-17, human reference build hg19) (Wang K. et al., 2010) to obtain the mutation effect, population allele frequency (AF), *etc*. After annotation, variants were selected based on the following criteria: 1. have a “PASS” value in the FILTER field; 2. have an AF < 5% in database gnomAD_genome_ALL; 3. exonic variants with the functional categories of “non-synonymous”, “frameshift_deletion”, “frameshift_insertion”, “nonframeshift_deletion”, “nonframeshift_ insertion”, “stopgain”, and “stoploss”; 4. not in dispensable genes as defined by (MacArthur D. G. et al., 2012) and (Rausell A. et al., 2020).

### Immunocytochemistry (ICC)

Cells were fixed in 4% PFA for 15 min and then permeabilized with 0.3% Triton X-100 in PBS for 10 min. Then, cells were blocked with 10% Normal Goat Serum for 1 h and incubated in the appropriate primary antibodies from mouse or rabbit: Sox2 (1:1000, Abcam, ab92494), Oct4 (1:250, Santa Cruz, Sc-5279), Nestin (1:2000–1:5000, R&D Systems, MAB1259), Pax6 (1:300, Covance, PRB-278P), β-III tubulin (TuJ1, 1:2000–1:5000, Covance, MMS-435P), Tau (1:500, Santa Cruz, Sc-5587), O4 (R&D Systems, MAB1326 1:500), GalC (Santa Cruz 1:250 sc-518055), and Glial fibrillary acidic protein (GFAP, 1:1000, Dako, G9269). Then, cells were washed three times for 5 min with 1X PBS and incubated 1 h in appropriate secondary antibody at 1:1000. The secondary antibodies used were red (594 Alexa Fluor, (ThermoFisher Scientific, Invitrogen, A-11032, A-11037) or green (488 Alexa Fluor ThermoFisher Scientific, Invitrogen, A-11001, A-11070) goat-anti-mouse antibodies or goat-anti-rabbit antibodies. Cells were visualized on fluorescent microscope.

### Media for Experimental Conditions

The protocols for all experimental media conditions, neurite outgrowth, and neurosphere assays are described in detail in our prior JOVE paper (Williams et al., 2018).

All experiments were conducted in 30 % expansion media (30 Exp) which was made by diluting Neural Expansion Media by 70% in a 1:1 ratio of DMEM/F12 and NB media. Primocin antibiotic (100ug/mL, InvivoGen ant-pm-1) was also added to the media

### Neurite Outgrowth Assay

50,000 NPCs were plated onto a 35 mm dish coated with 0.1 mg/mL PDL and 5ug/mL Fibronectin (Sigma Aldrich F1141) containing 1 mL of 30% Expansion media without or with drugs/growth factors. Vehicle, EF, or small molecule inhibitor/activator were added to media prior to cell plating. The EFs and concentrations used include: Nerve Growth Factor (NGF, Peprotech, AF-450-01) at 3, 10, 30 and 100 nM dissolved into 30% expansion media; PACAP (American Peptide/BACHEM, H-8430) at 1, 3, 10, 30 and 100 nM dissolved into 30% Expansion Media; and Serotonin (Sigma-Aldrich, H9523) at 30, 100, 300 ug/mL dissolved into 30% expansion media. The small molecule activator and inhibitor and concentrations were: SC79 (AKT activator, S7863) in DMSO diluted into 30% expansion media at concentration of 0.1, 0.3, 1, 3, 6 ug/mL and MK-2206 (AKT inhibitor, S1078) in DMSO diluted into 30% expansion media at concentrations of 0.3, 1, 3, 10, 30 and 100 nM. Vehicles were matched to whichever carrier was used to dissolve the drug or EF (for example for small molecule studies, vehicle was DMSO dissolved into 30% expansion media and volumes equivalent to the drug were added to each dish). Drugs and EFs were not replenished, and media was not changed in the 48-hour period of incubation and as such the initial EFs/molecules added in the start of the experiment remained for the full 48h period. Of note both MK-2206 and SC-79 act upon the pleckstrin homology domain of AKT to exert effects.

For each condition (control or EF or small molecule), 2-3 dishes were set up. NPCs were incubated at 37°C for 48 h and then fixed in 4% PFA. In each dish, the proportion of cells bearing neurites was counted blind. Neurites were defined as processes that extend from the cell body that are equal to or greater than 2 cell body diameters in length. For cells with more than one process, the longest process was assessed. Cells were counted directly on a phase contrast microscope at 32x. Two to four randomly chosen 1 cm rows were counted per dish. At least 250 cells were counted per dish and the proportion of cells with neurites was calculated in each dish.

### Neurosphere Migration Assay

Neurospheres were formed by dissociating confluent NPCs and plating 1 million cells into uncoated 35 mm dishes with 100 Exp. No small molecules or EFs were added during this formation step. The NPCs were then incubated at 37°C for 24-96 h (varied from line to line) to allow the aggregation of NPCs into neurospheres. Sphere size was assessed daily using a live-ruler on a phase-contrast microscope. When a majority of spheres reached an approximate diameter of 100 μm (± 20 μm), the migration assay was performed.

To coat plates, a Matrigel aliquot was dissolved into 6 mL of ice cold 30 Exp. Appropriate vehicles and EFs were added to the 30Exp/Matrigel mixture as desired. For migration the following EFs and small molecule inhibitors were used as noted above in the neurite section (PACAP, SC-79, MK-2206). Then, 1mL of the 30Exp/Matrix + EF mixture was added into the 6-well plate. Plates were then incubated for 30 min at 37°C. While plates were incubating, neurospheres were collected from the 35 mm dishes and placed into a 15 mL conical tube. The tube was centrifuged at 100g and the pelleted spheres were gently resuspended using a P1000 in 1-3 mL of pre-warmed 30 Exp (with appropriate concentration of EF or small molecule added). 200 μL of the sphere suspension was then placed into the 30 Exp Matrigel plates. Spheres were allowed to migrate for 48 hours and were fixed for 30 min in 4% PFA, washed, and stored in PBS + 0.05% Sodium Azide.

Neurospheres were imaged at 10X on a phase contrast microscope. At least 30 neurospheres across all dishes were imaged per condition. Spheres that were not in contact with another, that exhibited a contiguous migrating cell carpet and intact inner cell mass were imaged. Average migration was measured using ImageJ. To measure migration, the outer contour of a neurosphere was traced using the freehand line tool and the enclosed area was measured. Then, the inner cell mass area was measured. Migration was quantified by subtracting the inner cell mass area from the total neurosphere area. At least 20 neurospheres were analyzed per condition. Exclusion criteria are detailed in JOVE methods paper (Williams M. et al., 2018).

### Western Blot

To collect protein, confluent NPCs were dissociated, pelleted, and plated onto PDL+ Fibronectin coated plates with 1 mL 30% Exp. NPCs were plated at 1 million cells in 35 mm dishes and incubated for 48 h at 37°C. At 48 h, cells were treated with vehicle or drug for 20 or 30 minutes. Then, dishes were washed with ice-cold PBS 2 times. Cells were then lysed by adding 30 μL of ice-cold lysis buffer/dish and scraping with cell scraper. Lysate was then sonicated 2X for 1 min on ice and then samples were centrifuged at 4°C for 10 min. The supernatant was removed and saved while the pellet was discarded. Protein was aliquoted and stored at -80°C.

Aliquoted samples were thawed and 20 μg of protein was mixed with appropriate volumes of 4x NuPAGE LDS Sample buffer and 10x NuPAGE Sample Reducing Agent. All immunoblotting reagents were obtained from ThermoFisher Scientific. Samples were boiled, cooled, and loaded into wells on a 12% SDS-PAGE poly-acrylamide gel. The samples were run at 100 V for 1.5 h in an electrophoresis apparatus with 1X NuPAGE Mops SDS Running Buffer. Protein was then transferred to a PVDF membrane using a wet transfer apparatus with 1X NuPAGE Transfer buffer. The membranes were washed with Tris-Buffered Saline (TBS) with Tween20 (0.1%) (TBS-T) and blocked with 5% powdered milk in 0.1 %TBS-T. Then, membranes were probed for proteins with the following antibodies used at a 1:2000 concentration and incubated overnight at 4°C: S6 (2137 CellSignaling), P-S6 (2211, CellSignaling) as previously reported (Genestine M. et al., 2015; Yan Y. et al., 2013b; Mairet-Coello G. et al., 2009; Mairet-Coello G. et al., 2012; Tury A. et al., 2011). Total and Phospho-antibodies were run on different gels/ different halves of one gel to avoid stripping. GADPH was used at as a loading control and was probed with antibody (Meridian Life Sciences, H86045M) at a dilution of 1:10,000 for 1 h. After incubation in 1°, membrane was washed and appropriate HRP conjugated 2° antibody was applied at a 1:1000 concentration for all antibodies except GAPDH, for which 2° antibody was at 1:5000. Then, ECL was applied to membranes for 1 min and membrane was applied to medical grade X-ray film. Films were scanned into JPEGs and quantified on ImageJ. Band intensities for phospho- and total protein were normalized to signals for GAPDH. The GAPDH normalized Phospho-antibody intensity was divided by the GAPDH normalized total antibody to get a relative protein intensity.

### Phospho-Proteome

#### Protein Collection

As described above, protein was collected from NPCs plated on 0.1 mg/mL PDL+ 5μg/mL FN at a density of 1 million in 35 mm dishes for 48 hours. At least 1.5 mg of protein was collected per individual for p-proteomics. On average, 50-150 μg of protein was acquired from one dish of 1 million cells. Thus, to acquire 1.5 mg of protein, samples of protein were pooled from multiple experiments over time. For final samples, protein was collected from 3-4 experiments (of 3-5 dishes) across 3 different passages and two different neural inductions to ensure adequate sample and to account for variability. Samples were frozen at -80 after each collection and then ultimately thawed, pooled, and sonicated.

#### Proteomic Analysis

500 ug of each lysate from above were digested using standard FASTP protocol while 450 ug of each sample was labeled with TMT10 plex reagent (Thermo Scientific) according to manufacturer’s instructions and combined after labeling and dried (Mertins P. et al., 2013). TMT labeled and combined sample were desalted with SPEC C18 column and solubilized in 200 µL of buffer A (20 mm ammonium formate, pH 10) and separated on an Xbridge column (Waters; C18; 3.5 µm, 2.1 × 150 mm) using a linear gradient of 1% B min^−1^, then from 2% to 45% B (20 mm ammonium formate in 90% acetonitrile, pH 10) at a flow rate of 200 µL min^−1^ using Agilent HP1100. Fractions were collected at 1-min intervals and dried under vacuum. For total proteome analysis, 14 fractions (Fraction from 27 min to 40 min) were chosen. 5% of each fraction were desalted with stage tip and analyzed by nano-LC-MSMS. For phospho-proteomics, fractions were combined in concatenate style to make 6 fractions.

IMAC enrichment of phosphopeptides was adapted from Mertins et al (Mertins P. et al., 2013) with modifications. Ion-chelated IMAC beads were prepared from Ni-NTA Superflow agarose beads (Qiagen, MA). Nickel ion was stripped with 50 mM EDTA and iron was chelated by passing the beads through aqueous solution of 200 mM FeCl^3^ followed by three times of water wash and one time wash with binding buffer (40% acetonitrile, 1% formic acid). Combined HPH RP fractions were solubilized in binding buffer and incubated with IMAC beads for 1 hour. After three times of wash with binding buffer, the phosphopeptides were eluted with 2x beads volume of 500 mM potassium hydrogen phosphate, pH7.0 and the eluate was neutralized with 10% formic acid. The enriched phosphopeptides were further desalted by Empore 3M C18 (2215) StageTip^2^ prior to nanoLC-MS/MS analysis.

Nano-LC-MSMS was performed using a Dionex rapid-separation liquid chromatography system interfaced with a QExactive HF (Thermo Fisher Scientific). Samples were loaded onto a Acclaim PepMap 100 trap column (75 µm x 2 cm, ThermoFisher) and washed with Buffer A (0.1% trifluoroacetic acid) for 5 min with a flow rate of 5 µl/min. The trap was brought in-line with the nano analytical column (nanoEase, MZ peptide BEH C18, 130A, 1.7um, 75umx20cm, Waters) with flow rate of 300 nL/min with a multistep gradient. Mass spectrometry data were acquired using a data-dependent acquisition procedure with a cyclic series of a full scan acquired with a resolution of 120,000 followed by tandem mass spectrometry (MS/MS) scans (30% collision energy in the HCD cell) with resolution of 30,000 of the 20 most intense ions with dynamic exclusion duration of 20 s.

For both proteome and P-proteome studies, Maxquant identification and Perseus for quantitation were employed and an in-house R program that compounds variability at each level of the analysis (spectra to peptides, peptides to proteins, proteins per experimental group) was utilized to determine statistical significance. Specifically, significance for group comparisons used student T-tests with equal variance on both sites and Q values will be calculated using a 5% false discovery rate (FDR). Volcano plots were generated to illustrate individual peptides and P-peptides that are different between ASD cases and controls. Both statistically significant differences (P-value ≤ 0.05/number of observations after Bonferroni correction) and > 2-fold differences peptide amounts will be denoted. As such Initial data included over 9700 proteins, however, to appropriately adjust for multiple comparisons we only selected proteins which met the criteria of significance threshold greater than Log P of 5 for further analyses. This analysis was performed between: I-ASD vs Sib controls, 16p-Del vs Sib controls, and I-ASD vs 16pDel NPCs.

#### G-profiler Analysis

Our phosphor-proteomic data set, which included UniProt Identifiers and comparisons of fold changes relative to control (Sib), were submitted into the system. The specific tool utilized was g: GOSt which performs performs statistical enrichment analysis to interpret a user-provided gene list. The data provided users multiple sources of functional evidence including Gene Ontology (GO) terms, biological pathways, regulatory motifs of transcription factors and microRNAs, human disease annotations and protein-protein interactions. For analysis, g:Profiler allows users to set preferences on statistical domain scope, significance threshold, and user threshold. Briefly, for the statistical domain scope, the program was tasked to look through all known genes, allowing for calculation of statistical significance considering all genes of the human genome in the Ensembl database. Next, for the significance threshold the g:SCS algorithm was used, which sets an experiment-wide threshold of p = 0.05. Finally, the user defined p-value threshold allows for further filtering of results; this threshold was also set to p = 0.05. As a readout of the results, the program looks at Gene Ontology (GO), which is a bioinformatics initiative that defines biological functions and their relationship to one another. The three major subontologies are Molecular Functions (MF), gene products functions, Biological Process (BP), which describes biological processes the gene product participates in, and Cellular Component (CC), describes which part of the cell the gene product is physically located.

#### Ingenuity Pathway Analysis (IPA)

Our phospho and total proteomic data set, which included UniProt Identifiers, P-values, Q values and log2 ratios of comparisons of fold changes of identified phosphorylated proteins relative to control (Sib), were submitted into the IPA system for core analysis. The pathways and functional networks presented were all generated through the use of IPA QIAGEN Inc., https://www.qiagenbioinformatics.com/products/ingenuity-pathway-analysis/). The core analysis generated molecular networks according to biological as well as molecular functions including canonical pathways, disease-based functions, and upstream regulatory analysis for the discovery of strongly differential molecules in our dataset. Both direct and indirect relationship between molecules based on experimentally observed data, and data sources were considered in human databases in the Ingenuity Knowledge Base. Right-tailed Fisher’s exact test was used to determine the probability that biological functions and/or diseases were over-represented in the protein dataset. Z-scores compiled as a statistical result of differential protein expression according to the fold changes was also compiled.

### Compilation of Data and Statistical Analyses

Data acquired from the idiopathic autism individuals were compared to data acquired from unaffected Sibs. Data was compiled and organized in several ways. Comparisons were made within Family (Sib-1 vs ASD-1), across families (Sib-1 vs ASD-2), or as an average of all Sib (3 patients, all clones) compared to the average data from all idiopathic ASD (3 patients, all clones). For every neurite experiment, 2-3 dishes were set up per condition. For every migration condition, 20-30 spheres were evaluated per condition. Dishes were kept as separate N values and were not collapsed into a single mean per experiment. For example, when compiling data for Sib-1 in control condition, all neurite dishes from every clone were averaged together to get a mean neurite value for this individual. For spheres, each neurosphere was considered a separate statistical point. Migration values from each sphere derived from multiple clones were averaged together to acquire an average migration value for each patient. Spheres were also analyzed by clone. For the 16p11.2, data was compared to composite averages of the Sibs (dishes from all 3 Sib patients and all clones) or averages of the NIMH controls (dishes from both patients, all neural inductions). Assuming statistical normality, in comparisons that include only two groups, unpaired t-test was used to test differences in affected vs unaffected samples. Simple unpaired or paired t-tests were run in Microsoft Excel. If normality assumption was not satisfied, the non-parametric Wilcoxon test was used in Graphpad Prism 8. The presence of multiple ASD subtypes and the use of EFs leads to instances where multiple group comparison is necessary. For such cases, in normal data, analysis of variance or ANOVA (one or two way) was used to detect statistical significance. To reduce type one error, P-values for the ANOVAs were calculated using Tukey correction in GraphPad Prism 8. If normality assumption did not hold, non-parametric Kruskal-Wallis test was applied for multiple comparisons.

## RESULTS

### Patient Cohort, Rigor and Reproducibility, and Study Design

The I-ASD cohort was selected from the previously reported New Jersey Language and Autism Genetics Study. From 3 separate families, iPSCs were generated from 3 male probands with ASD and their sex-matched unaffected siblings (Sib) served as controls (see methods).

For comparison, we selected a genetically-defined ASD cohort with 16p11.2 deletion CNV. The CNV deletes 28 genes, increases autism risk by ∼20-fold and is one of the most common ASD-associated mutations (Weiss L. A. et al., 2008; Niarchou M. et al., 2019). This cohort consists of 3 individuals, 2 males (16pDel M-1, 16pDel M-2) and 1 female (16pDel F). For sex-matched controls, we selected 3 iPSC lines from the NIH Regenerative Medicine Program (NIH).

We analyzed these 12 individuals (six I-ASD-Sib and six 16pDel-NIH) extensively for developmental phenotypes utilizing 2-5 clones/individual (Except for NIH) and 2-5 independent neural inductions to ensure extensive rigor and reproducibility (see methods and SI Table 1).

### Whole Genome Sequencing

WGS was conducted on I-ASD dataset and demonstrated that no individuals harbored the 16p11.2 deletion (SI Table 2). Moreover, no I-ASD individual had rare non-synonymous protein coding variants in the 28 genes deleted in 16p11.2. Finally, there are no rare protein-coding non-synonymous genetic variants shared between all 3 I-ASD individuals. These data are consistent with the two datasets being genetically distinct with no common or shared genetic drivers.

### I-ASD and 16pDel have Common Defects in Neurite Outgrowth and Cell Migration

To study an ASD-relevant developmental process, we first assessed neurite outgrowth in I-ASD NPCs at 48h (Fig. 1A). Given the genetic heterogeneity of the I-ASD cohort, we expected to find personalized phenotypes. Yet, surprisingly, we found that all 3 I-ASD NPCs had a significantly lower percentage of neurites than their respective Sibs (Fig. 1B, 50-60% reduction, p<0.001 for all families). To understand how I-ASD NPCs compared to genetically unrelated neurotypical individuals, we also compared I-ASD NPCs to Sibs from other families (ex: I-ASD-1 to Sib-2). On average, all I-ASD NPCs had reduced neurites compared to all Sib NPCs.

Given the common neurite impairment in all I-ASD NPCs, we then extended our studies to another early developmental process, NPC migration, which occurs concurrently with neurite outgrowth and is similarly regulated during development (Prem S. et al., 2020). For these studies, we plated neurospheres and at 48h and measured the area covered by migrating NPCs (Fig. 1C). Remarkably, we found that all 3 I-ASD patients exhibited decreased migration (∼30-50%) compared to their Sib (p<0.001; Fig. 1D) as well as unrelated sibs. Thus, despite genetic heterogeneity both neurite outgrowth and cell migration are impaired in all 3 I-ASD NPCs.

This remarkable convergence prompted us to examine the 16pDel ASD cohort for these phenotypes. Fascinatingly, 16pDel NPCs have a lower percentage of neurites (7.4%) than both the Sib (11.3%, p<0.001) and NIH controls (12.7% p<0.0001). This indicates that defects in neurite outgrowth are a common feature across two genetically distinct subtypes of autism. Interestingly, 16pDel NPCs have a higher percent of neurites than I-ASD (7.4% vs 5.1% p<0.01) indicating potential subgroup differences. There was no difference in neurite extension between Sib NPCs (11.3%) and NIH NPCs (12.7%) (p=0.1).

Lastly, we assessed migration in 16pDel NPCs. Once again, like I-ASD NPCs, all 3 16pDel NPCs have impaired migration when compared to Sib and NIH NPCs (Fig. 1F). There were no differences in migration between 16pDel and I-ASD or Sib NPCs and NIH NPCs.

In summary, we have found that two genetically distinct ASD cohorts have surprisingly common dysregulation in two early neurodevelopmental processes, neurite outgrowth and cell migration. Given the marked heterogeneity of ASD, these results are striking and potentially suggest that common mechanisms regulate these neurodevelopmental defects.

### I-ASD and 16pDel NPCs Have Subtype-Specific Responses to Extracellular Factors (EFs)

Few iPSC studies have explored the roles of EFs in regulating human neural cell development. We postulated that stimulating NPCs with EFs could reveal defects not apparent in control conditions and might help identify underlying downstream signaling defects. Thus, we stimulated our NPCs with NGF, PACAP, and 5-HT which regulate neurite outgrowth or cell migration in rodent studies. (Maisonpierre P. C. et al., 1990; Hanswijk S. I. et al., 2020; Dicicco-Bloom E. et al., 1998; Lu N. et al., 1997).

We first tested EFs on the I-ASD cohort. Under PACAP, NGF, and 5-HT stimulation, Sib NPCs from all 3 families had significant increases (45-75%, p<0.001) in neurite outgrowth (Fig. 2A, B, and S1A). In contrast, all 3 I-ASD NPCs failed to respond to the three EFs (Figs. 2A, B, and S1A). The lack of response was not due to differential sensitivities to or inappropriate doses of EF, as dose response analyses indicate Sibs respond to PACAP and NGF at multiple doses while ASD had no response at any physiological dose (Fig S1B, C).

**Figure 2:**
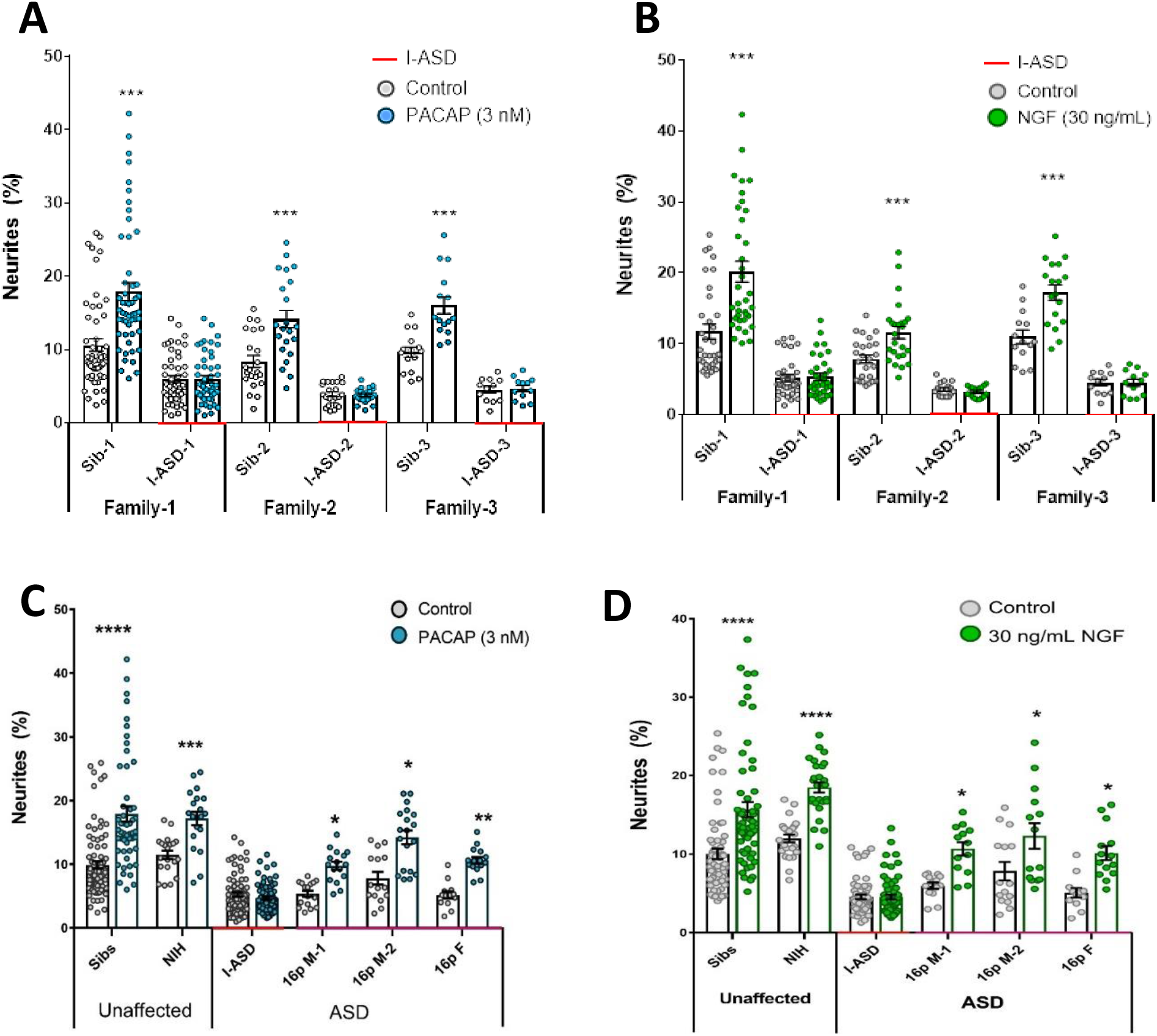
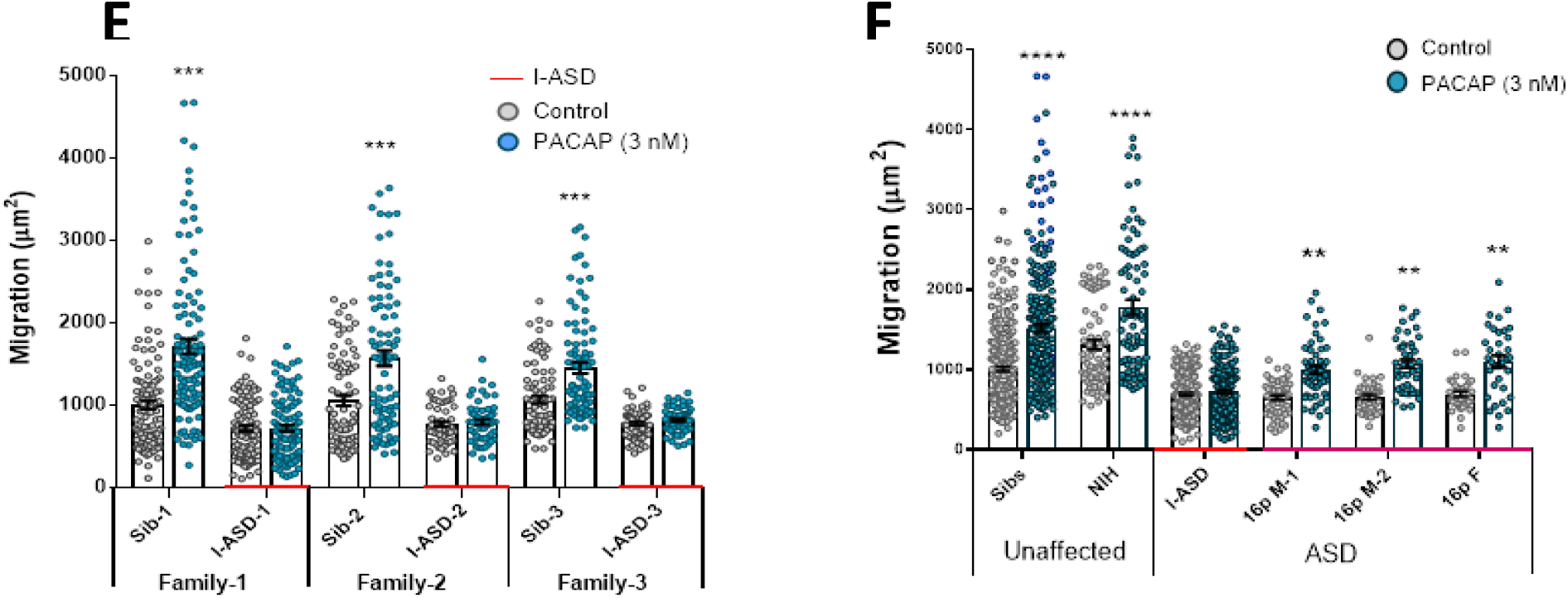
ASD Subtype Specific EF responses in I-ASD and 16pDel NPCs. A and B) 3nM PACAP (A) and 30ng/mL NGF (B) increases neurite outgrowth in all Sibs but fails to stimulate neurite outgrowth in all I-ASD NPCs. C and D) 3nM PACAP (C) and 30ng/mL NGF (D) stimulates neurite outgrowth in Sibs, NIH, and all 3 16pDel NPC but not in I-ASD NPCs. E) 3nM PACAP increases cell migration in all Sib NPCs but fails to stimulate migration in all I-ASD NPCs. F) 3 nM PACAP stimulates migration in Sibs, NIH, I-ASD, and all 3 16pDel patient NPCs but does not change migration in I-ASD NPCs. 2-Way ANOVA, p<0.01 for all unaffected vs affected comparisons.

We also applied EFs, specifically PACAP, a well-known regulator of migration in rodent models (Falluel-Morel A. et al., 2005), to our migration assay. Like our neurite results, increased cell migration in Sibs was observed but there was no effect in all 3 I-ASD NPCs (Fig. 2E).

Next, we extended EF studies to the 16pDel cohort. Surprisingly, unlike I-ASD NPCs, all three 16pDel NPCs respond to EFs with increases in neurite outgrowth and migration (Fig 2C, D, E, F, S1D). As expected, like Sib controls, NIH NPCs also exhibit increases in neurite outgrowth and migration by EF stimulation (Fig 2C, D, E, F, S1D).

In summary, while our two ASD cohorts have common impairments in neurite outgrowth and migration, EF experiments uncovered subtype specific response in I-ASD vs 16pDel. Thus, subtype-specific defects can be present alongside common neurobiological phenotypes.

### Phospho-Proteomic Analysis Reveals Dysregulated Signaling in Both ASD Cohorts

Signaling pathways are central to the regulation of neurodevelopmental processes and signaling dysregulation has been implicated in NDD pathogenesis (Lipton J. O. et al., 2014; Takei N. et al., 2014; Kelley D. J. et al., 2008; Pucilowska J. et al., 2012; Kwan V. et al., 2016; Wang L. et al., 2017; Waite K. et al., 2011). Given the neurodevelopmental phenotypes present in all our ASD patients and the impaired EF responses in I-ASD, we postulated that dysregulated signaling pathways could be an underlying mechanism. Thus, we conducted unbiased proteomic and p-proteomic studies comparing Sib and I-ASD, Sib and 16pDel, and I-ASD and 16pDel.

We first analyzed the proteome and found surprisingly few significant changes relative to Sib in the I-ASD and 16pDel datasets. Initial data included over 9700 proteins, however, to adjust for multiple comparisons we only selected proteins which met significance threshold of LogP >5 for further analyses. Given these parameters, I-ASD proteome had 9 changes compared to Sib whereas 16pDel had 48 changes compared to Sib. There were only 3 proteins (TAPT-1, GGA-1, S100A11) in common between I-ASD and 16pDel proteomes (Fig 3A).

**Figure 3:**
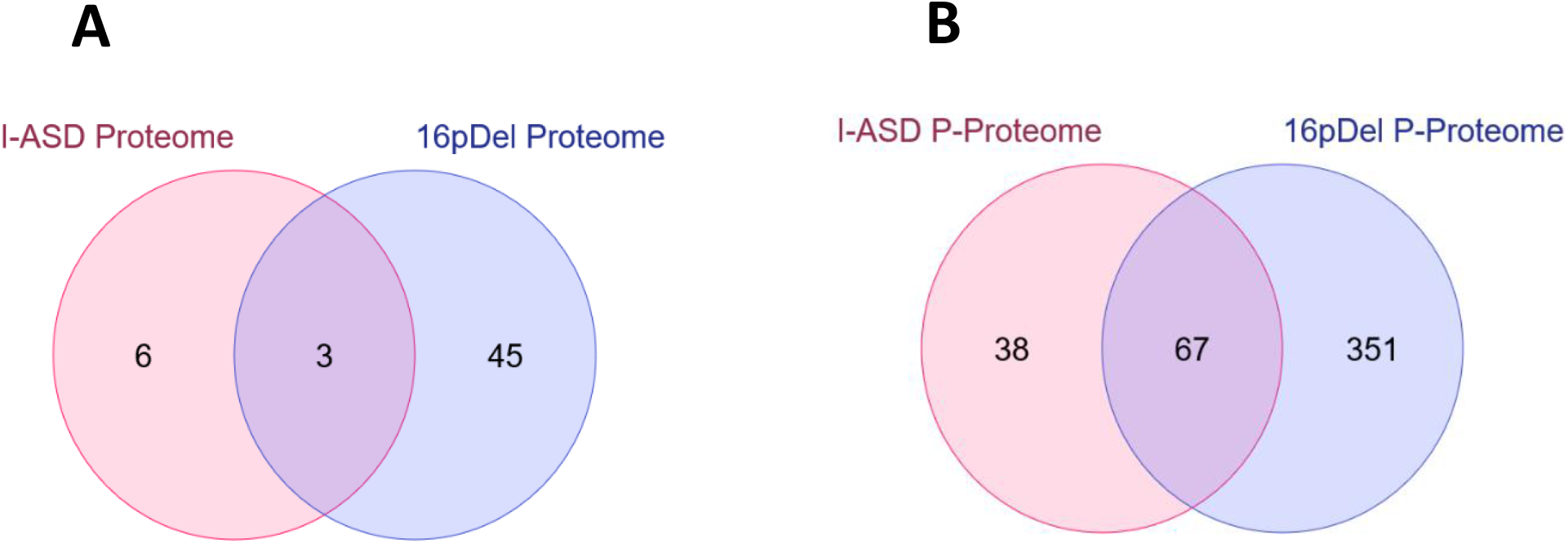
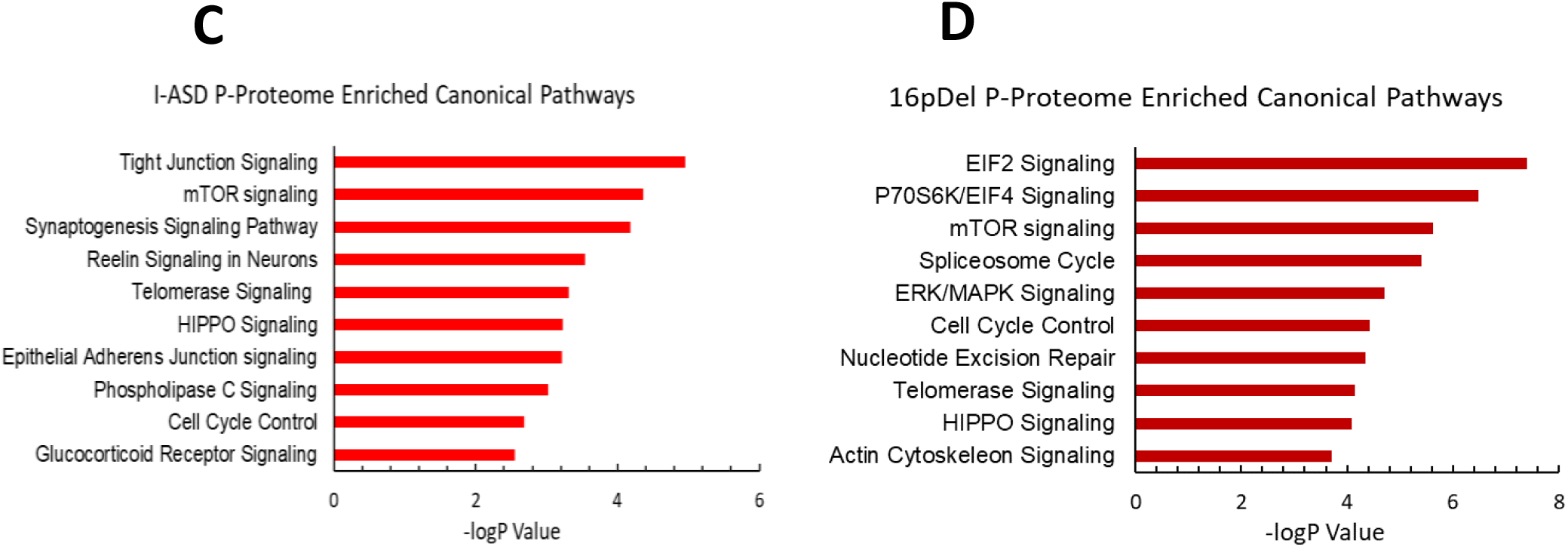
Phospho-proteome analysis of I-ASD and 16pDel Data: A and B) Venn Diagram illustrates Proteomic (A) and P-Proteomic (B) changes in both I-ASD and 16pDel NPCs and their overlap. C and D) IPA analysis of I-ASD (C) and 16pDel (D) p-proteome data identifies the mTOR pathway as likely disrupted.

In striking contrast, many more changes were observed in the p-proteome analysis: I-ASD NPCs had 105 differentially phosphorylated proteins compared to Sib with 149 unique phosphorylation sites.

Further, 16pDel NPCs had 419 differentially phosphorylated proteins with 916 unique phosphorylation sites. Surprisingly, given the absence of genetic overlap between the cohorts, there is considerable overlap between I-ASD and 16pDel p-proteome results with 67 p-proteins (67/105) and 107 phosphorylation sites (107/149) shared between I-ASD and 16pDel (Fig 3B). The large differences in the p-proteome along with minimal changes in the proteome suggests that phosphorylation and signaling could be dysregulated in both forms of ASD. Moreover, the p-proteome overlap between I-ASD and 16pDel NPCs suggests that different genetic mutations could converge on common signaling pathways.

To further assess p-proteome data, bioinformatic analyses were performed. To begin to understand the biological processes that could be dysregulated, G-profiler analysis was conducted (Raudvere U. et al., 2019; Reimand J. et al., 2019). For both I-ASD and 16pDel p-proteome, alterations in cytoskeleton, cell cycle, and developmental processes were identified, correlating well with our defined cellular phenotypes. To identify dysregulated signaling pathways, Qiagen Ingenuity Pathway Analysis (IPA) was utilized (Kramer A. et al., 2014; Schubert K. O. et al., 2015). IPA canonical analysis of I-ASD p-proteome determined the top three enriched modules are tight junction signaling, mTOR signaling, and synaptogenesis signaling (Fig 3C, S2A). Analysis of the 16pDel p-proteome reveals the top three are EIF2 signaling, regulation of E1F4 and p70S6K signaling, and mTOR signaling (Fig 3D, S2B). In striking contrast, IPA analysis of total proteome data even with threshold Log P >3) did not find any mTOR enrichment in either I-ASD or 16pDel NPCs (Fig S2C, D). Thus, the multi-component mTOR pathway which includes molecules like AKT and activation of p70S6K and ultimately phosphorylation of S6, is a common point of convergence between I-ASD and 16pDel. Indeed, IPA expression network analysis reveals strongly altered p-S6 as a central node of convergence in both I-ASD and 16pDel networks along with other mTOR members such as p-AKT (Fig S2E, F). Thus, despite the two datasets being genetically distinct, both I-ASD and 16pDel share many p-proteome changes with mTOR signaling and convergence onto RPS6 being common between them.

### mTOR Abnormalities are Found in all 6 ASD NPCs

To further assess the mTOR pathway, western blot studies were conducted to define p-S6 levels in 48h NPC cultures. Remarkably, p-S6 dysregulation was found in all 6 ASD NPCs from both cohorts (Fig 4). Further, less significant differences in upstream p-AKT levels were also noted in all ASD NPCs (Figure S3A-S3H). Importantly, total S6 and AKT protein levels were not different between any of the NPCs. While p-proteome revealed p-S6 as a common dysregulated node amongst all ASD individuals, our western studies revealed two distinct groups of p-S6 dysregulation. In one group, p-S6 levels were reduced relative to controls whereas in the other p-S6 levels were elevated, suggesting two distinct “signaling subtypes” as described below.

**Figure 4:**
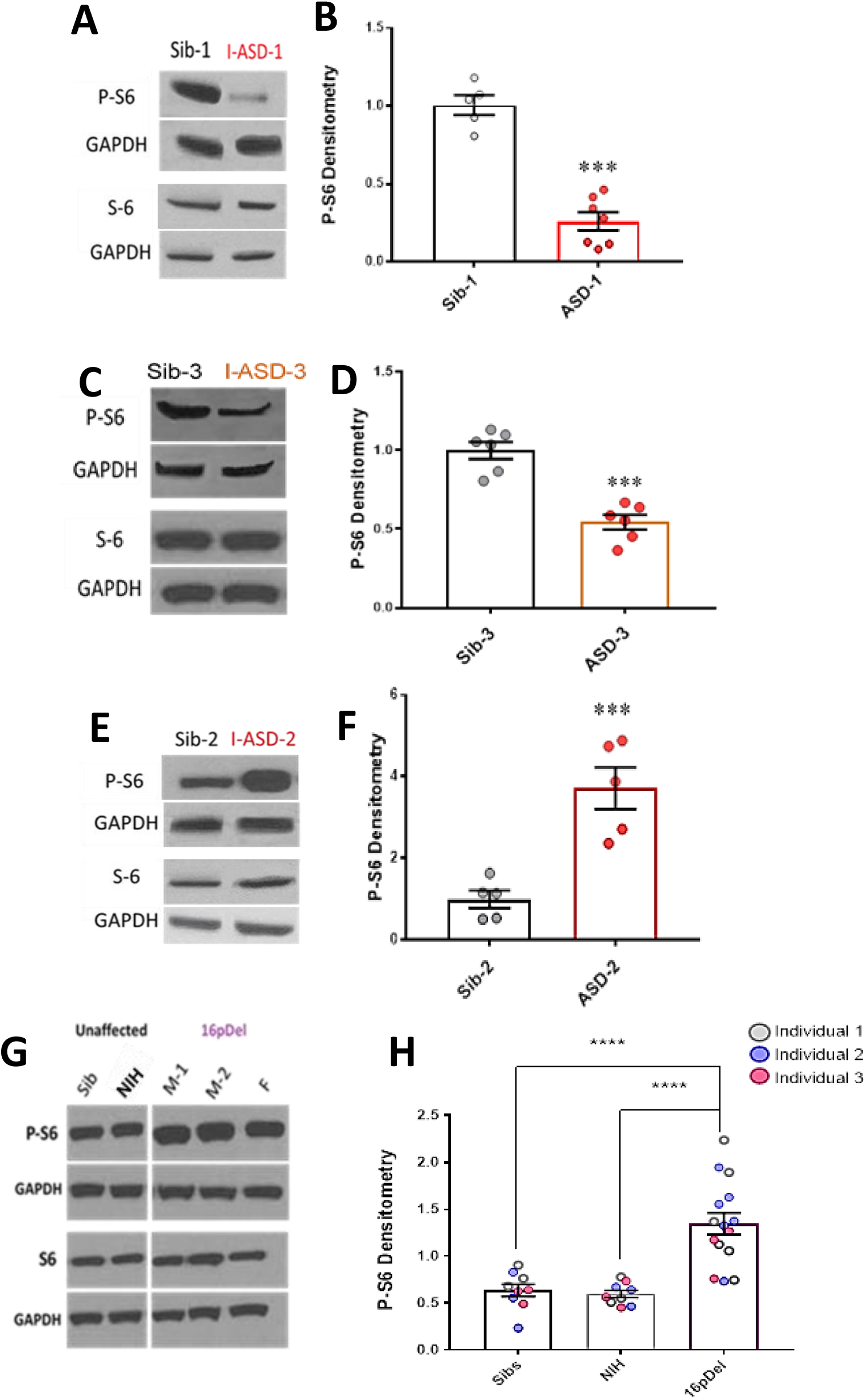
S6 and P-S6 western analysis for I-ASD and 16pDel NPCs. All images show with matched GAPDH loading control. Graphs show densitometry quantifications of normalized P-S6 (P-S6/GAPDH) divided by normalized S6 (S6/GAPDH). Student’s T-test for all I-ASD comparisons. A-D) I-ASD-1 and -3 representative western blots showing reduced P-S6 but similar S6 and GAPDH in I-ASD-1 compared to Sib-1 (A) and in I-ASD-3 vs. Sib-3 (C). Graphs (B and D): reduced P-S6/S6 in I-ASD-1 vs. Sib-1 (p<0.0001) (B) and in I-ASD-3 vs Sib-3 (D) (p<0.0001). E) I-ASD-2 representative western blot showing increased P-S6 but similar S6 in I-ASD-2 compared to Sib-2. F) Graph: Elevated P-S6/S6 in I-ASD-2 compared to Sib-2 (p<0.001). G) Western blot comparing a representative Sib, NIH control and each 16pDel patient (M-1, M-2, F) showing increased P-S6 in all 16pDel NPCs compared to both NIH and Sib with similar total S6 and GAPDH. H) Graph showing increased P-S6 in 16pDel compared to both Sibs and NIH (p<0.001 1-way ANOVA).

In the I-ASD cohort, Family 1 (I-ASD-1) had a 75% reduction in p-S6 compared to Sib-1 (Fig 4A, B) while I-ASD-3 from Family 3 had a 45% reduction in p-S6 compared to Sib-3 (Fig 4C, D p<0.001). Surprisingly, in contrast, I-ASD-2 from Family 2 had elevated levels of p-S6 (Fig 4 E, F 145% increase, p<0.001) when compared to Sib-2. The abnormal p-S6 levels of each ASD NPC were also significant when compared to the mean of the 3 Sibs. (Fig S3I). Likewise, p-AKT data followed the same pattern but with a smaller effect (Fig S3 A-H). In the 16pDel group, all 3 16pDel patient NPCs had higher p-S6 levels compared to both Sib and NIH NPCs (M1: ∼170% p<0.0001, M2: ∼130% p<0.001, F: ∼80% p<0.01 increase, Fig. 4G, H). There were no statistical differences in p-S6 levels between Sib and NIH NPCs (p=0.99). All 16pDel NPCs also displayed higher p-AKT levels compared to control (Fig S3 G, H).

Thus, western studies validate the p-proteome results in our 6 ASD patients and confirm changes in both p-S6 and p-AKT, which can be divided into low mTOR and high mTOR groups. Ultimately, the most striking finding is the consistent mTOR signaling differences across two genetically distinct groups of autism, a finding not previously described

### mTOR Pathway Modulation Rescues and Recapitulates Common Neurodevelopmental Phenotypes in Low mTOR NPCs

To determine whether the mTOR differences described above drive the developmental abnormalities in our cohorts, we conducted gain and loss of function studies on all I-ASD and both male 16pDel NPCs. As there we no small molecule activator and inhibitor pairs for mTOR and S6 when experiments were being conducted, we employed an AKT activator (SC-79) and AKT inhibitor (MK-2206), which have been used extensively in other systems to modulate mTOR (Jo H. et al., 2012; Takizawa S. et al., 2012; Hirai H. et al., 2010). Notably, AKT phosphorylation leads to mTOR pathway activation and S6 phosphorylation.

We first examined our low mTOR cohort comprised of I-ASD-1 and I-ASD-3. Focusing first on Family-1, addition of the SC-79 activator (2µg/mL) to I-ASD-1 NPCs increased p-S6 and p-AKT levels to that of Sib (Fig. 5A, B, S4A, B). Importantly, SC-79 at this dose had no effect on p-S6 or p-AKT levels in Sib, nor on the levels of total S6 or AKT in either Sib or I-ASD. We next tested whether rescuing P-S6 with SC-79 affected neurite outgrowth or migration. Indeed, SC-79 increased the percentage of neurites (Fig 5C) and migration in I-ASD NPCs (Fig 5D) without changing these parameters in Sib. Thus, we could rescue the neurite and migration defects in I-ASD-1 by modulating mTOR signaling and increasing p-S6 levels.

**Figure 5:**
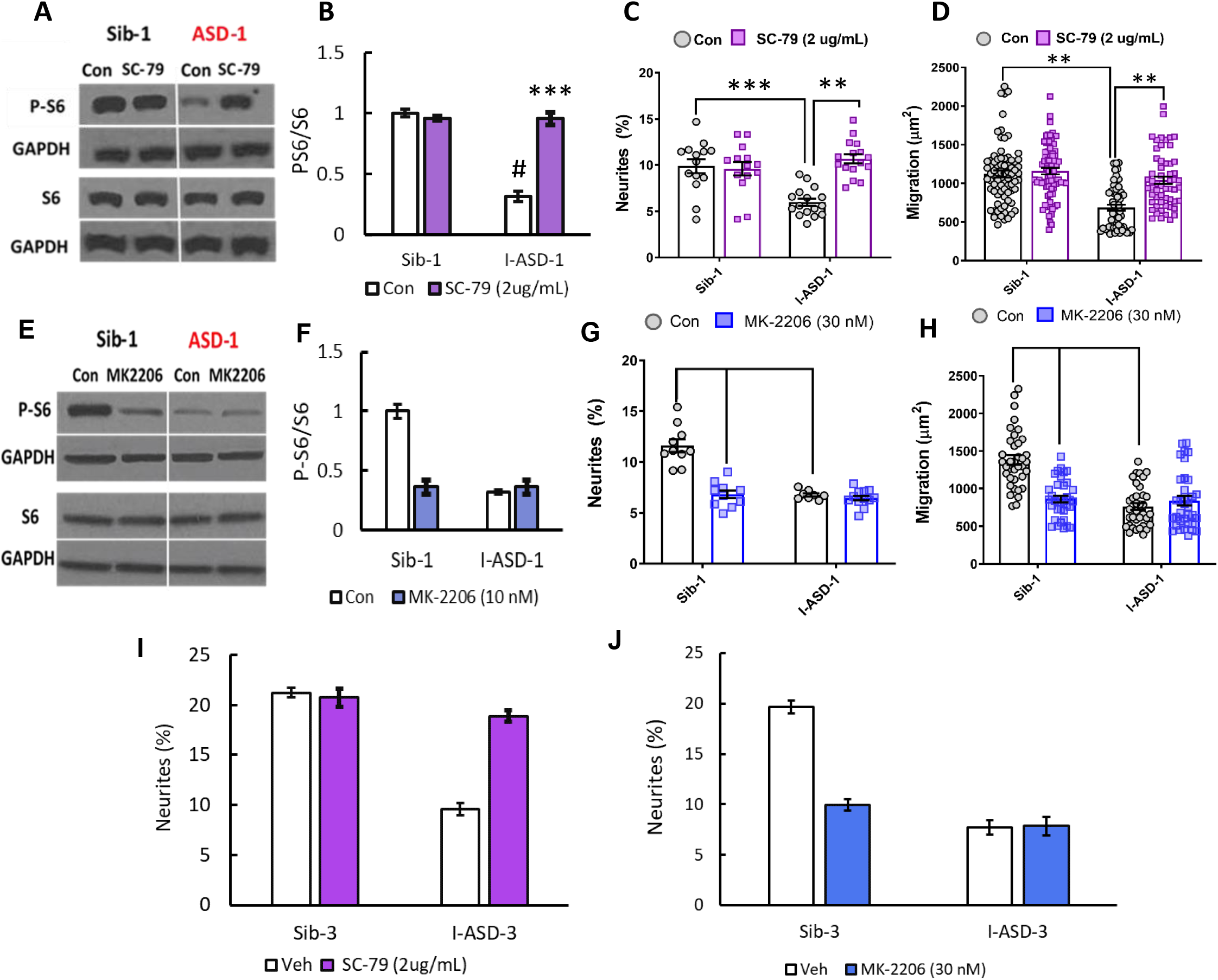
Effects of mTOR pathway manipulation in low mTOR cohort: A) Representative western blot: treatment of I-ASD-1 NPCs with SC-79 increases P-S6 levels in I-ASD-1 but not Sib-1 with no changes in total S6. B) Graph: SC-79 vs Veh: increases P-S6/S6 levels in I-ASD-1 but not in Sib-1. C and D) SC-79 treatment rescues both neurite outgrowth (C) and migration (D) in low mTOR I-ASD-1 NPCs without affecting Sib-1 NPC neurites. D) SC-79 treatment rescues migration in low mTOR I-ASD-1 NPCs without affecting Sib-1 NPC migration. E) Representative western blot: MK-2206 treatment of I-ASD-1 NPCs decreases P-S6 in Sib-1 but not I-ASD-1 with no changes in total S6. F) Graph: MK-2206 vs Veh: decreases PS6 in Sib-1 but not I-ASD-1. (G and H) MK-2206 treatment reduces neurite outgrowth (G) in Sib-1 to the level of I-ASD-1 without affecting I-ASD-1 neurite outgrowth. H) MK-2206 treatment reduces migration in Sib-1 to the level of I-ASD-1 without affecting I-ASD-1. I) SC-79 treatment of low mTOR I-ASD-3 increases neurite outgrowth to the level of Sib-3. J) Treatment of Sib-3 NPCs with MK-2206 reduces neurite outgrowth to the level of I-ASD-3. 2-way ANOVA for all analyses, p<0.01 for all labeled comparisons.

Of course, if mTOR pathway deficiency is truly contributing to ASD defects in neurite outgrowth and migration, then inhibitor MK-2206, should diminish neurite outgrowth and impair migration in control Sib-1 NPCs. First, by using MK-2206 (30nM) we were able to reduce p-S6 and p-AKT levels in Sib down to that of I-ASD-1 (Fig 5E, F, S4). MK-2206 exposure also reduced the percentage of neurites (Fig. 5G) and migration (Fig 5H) of Sib-1 NPCs to that of I-ASD-1 NPCs thereby reproducing both the ASD neurite and migration defects.

Lastly, we tested whether modulating p-S6 in I-ASD-3, the other “low mTOR” individual, also led to similar results. The addition of SC-79 activator to I-ASD-3 NPCs led to an increase in neurite outgrowth that paralleled Sib-3 NPCs (Fig. 5I). Likewise, inhibiting p-S6 with MK-2206 in Sib-3 led to a reduction in neurite outgrowth that mimics the phenotype seen in I-ASD-3 (Fig. 5J). Thus, in both “low-mTOR” ASD patients, we find that increasing mTOR pathway activity (via SC-79) can rescue ASD defects while reducing mTOR pathway activity (via MK-2206) in the Sibs reproduces autism phenotypes.

### mTOR Pathway Modulation Rescues and Recapitulates Common Neurodevelopmental Phenotypes in High mTOR NPCs

Given that increasing p-S6 in our low mTOR cohort led to rescue of ASD phenotypes, we next sought to determine whether decreasing p-S6 in high mTOR ASD NPCs (ASD-2 & 16pDel) could similarly rescue ASD phenotypes. Starting with I-ASD-2, addition of MK-2206 inhibitor reduced the levels of p-S6 to that of Sib without changing total S6 levels (Fig. 6A, B). This increased both neurite outgrowth and migration paralleling levels seen in Sib, thereby rescuing the phenotypes (Fig. 6C, D). Significantly, Sib-2 had no response to MK-2206 at this dose.

**Figure 6:**
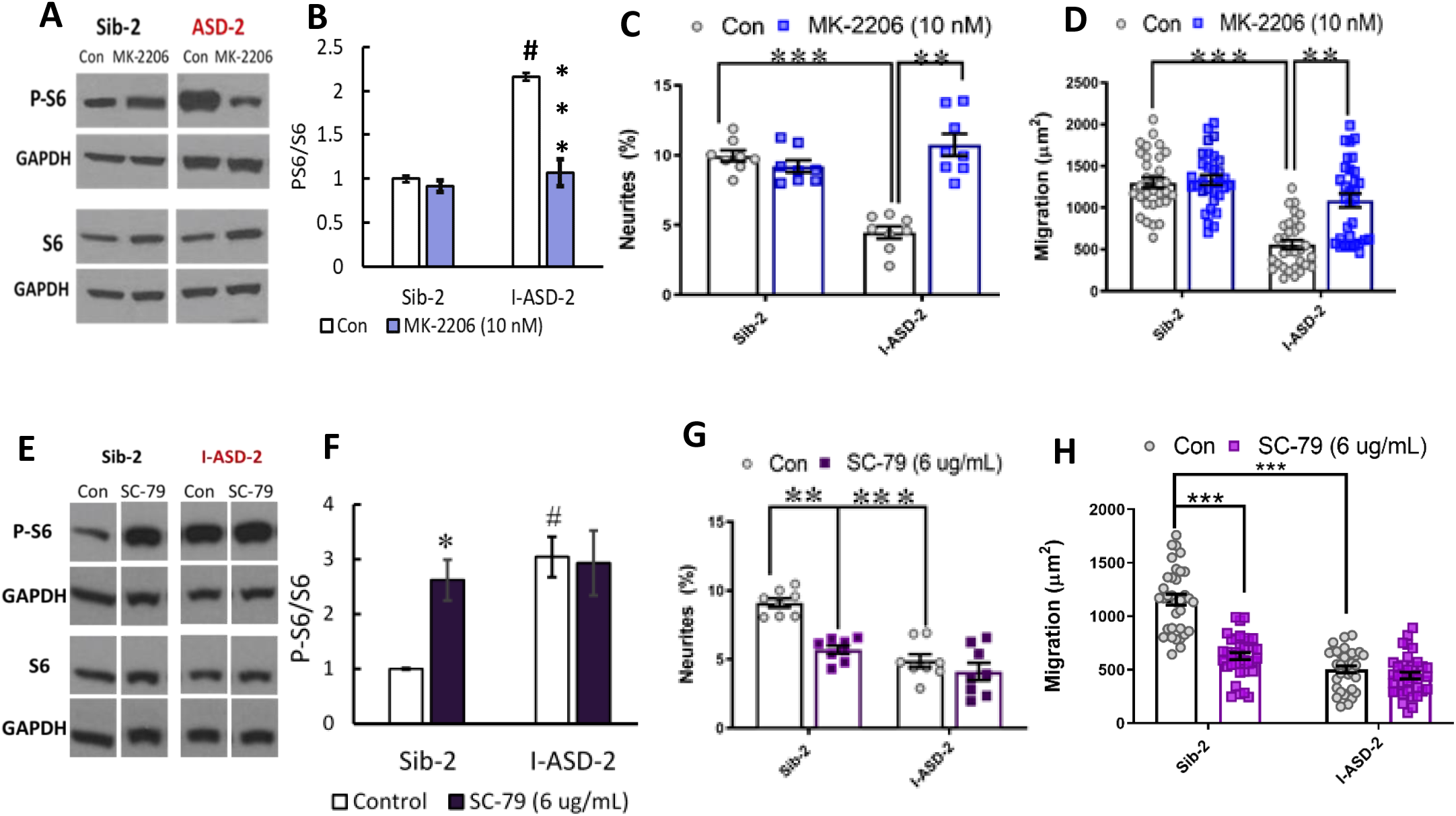
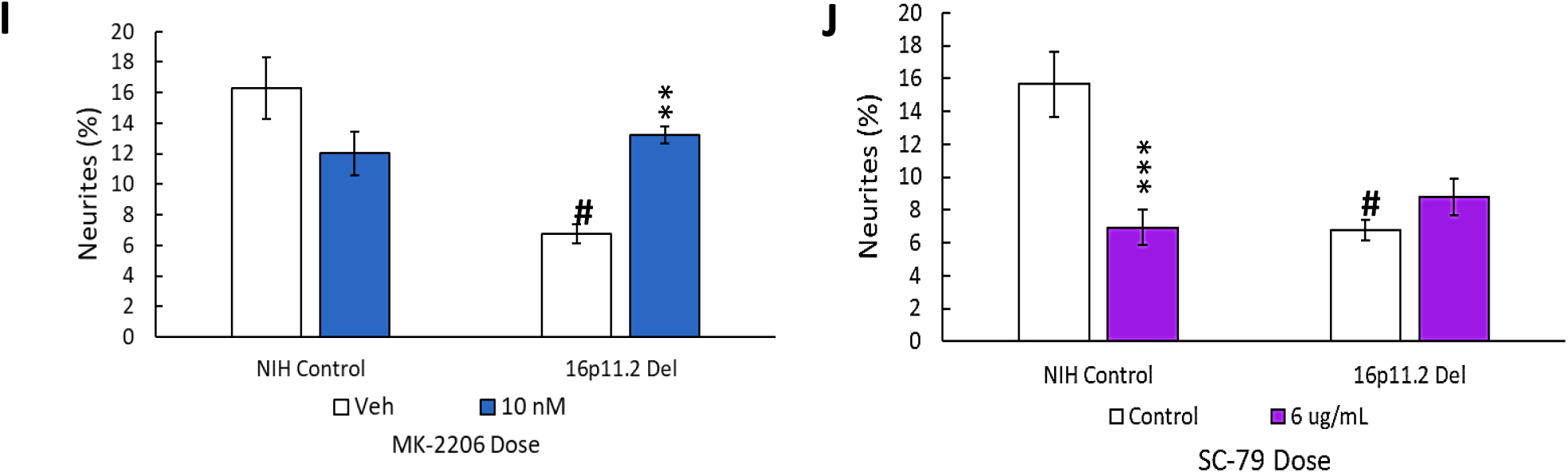
Effects of mTOR pathway manipulation on high mTOR cohort. A) Representative western: MK- 2206 treatment of I-ASD Family-2 NPCs leads to reduction of P-S6 in I-ASD-2 to Sib-2 levels without changing total S6. B) Graph: MK-2206 vs Veh - quantification of PS6/S6 western blots showing decreased PS6 in I-ASD-2 but no change in Sib-2. C and D) MK-2206 rescues neurite outgrowth (C) and migration (D) in high mTOR I-ASD-2 without changing Sib-2 neurites or migration. E) Representative western: SC-79 treatment of I-ASD Family-2 NPCs increases P-S6 in Sib-2 to the level of I-ASD-2 without changing total S6. F) Quantification of Veh vs.SC-79 westerns showing increased PS6 in Sib-2 but no change in I-ASD-2. G and H) Treatmentof Sib-2 with SC-79 diminished neurite outgrowth (G) and migration (H) to the level of I-ASD-2. I) In high mTOR 16pDel NPCs, MK-2206 treatment rescues neurites without significantly affecting NIH control NPCs. J) Treatment of NIH NPCs with SC-79 diminishes neurite outgrowth to 16pDel levels without affecting 16pDel NPCs. For all comparisons p <0.01 (2-way ANOVA).

We next determined if elevating mTOR signaling using SC-79 could reproduce the I-ASD-2 defects in the Sib. SC-79 increased p-S6 levels in Sib-2 NPCs to that of I-ASD-2 (Fig. 6E, F), which remarkably led to neurite and migration impairments that mimic those seen in I-ASD-2 (Fig 6 G, H).

Finally, we conducted gain and loss of function studies in the two male 16pDel NPC and NIH NPCs. Like I-ASD-2, the 16pDel NPCs have reduced neurite outgrowth and migration in conjunction with elevated p-S6. Reducing p-S6 levels with inhibitor (MK-2206) rescued the 16pDel phenotypes as evidenced by increased neurite outgrowth (Fig 6I). Conversely, increasing p-S6 with activator (SC-79) in NIH NPCs reproduced the neurite defects noted in the 16pDel NPCs (Fig 6J). In summary, we find that both over- and under-activation of mTOR signaling leads to a common reduction in neurite outgrowth and migration. Since these phenotypes could be rescued or reproduced in both low and high mTOR NPCs by manipulating mTOR signaling, they indicate the need for tight regulation of this pathway for normal neural development.

### Altering the mTOR Signaling Milieu Establishes EF Responsiveness in I-ASD NPCs

The above results indicate that mTOR signaling dysregulation is responsible for the shared neuro-developmental phenotypes. The I-ASD NPCs also do not respond to EFs (PACAP, NGF, and 5-HT) which in Sib controls stimulate neurite outgrowth and migration. To further investigate if this failure to respond to EFs is due to signaling defects, we treated the I-ASD NPCs with “subthreshold” amounts of mTOR agonist and inhibitor which did not maximally increase p-S6 or elicit neurite outgrowth on its own. To do so, we performed parallel dose response studies and selected drug doses that did not elicit neurite outgrowth on their own but did elicit a non-maximal p-S6 increase (data not shown). For Family-1, agonist SC-79 (0.1 µg/mL) did not stimulate neurite outgrowth in I-ASD-1 NPCs (Fig. 7A). However, when combined with PACAP, NGF, and 5-HT, I-ASD-1 NPCs now responded to all 3 EFs (Fig. 7A). This effect was specific to I-ASD-1 NPCs as addition of sub-threshold SC-79 to Sib NPCs did not heighten EF response (Fig 7B). Thus, by using sub-threshold doses of mTOR activator, we could facilitate EFs signaling in I-ASD-1 NPCs.

**Figure 7:**
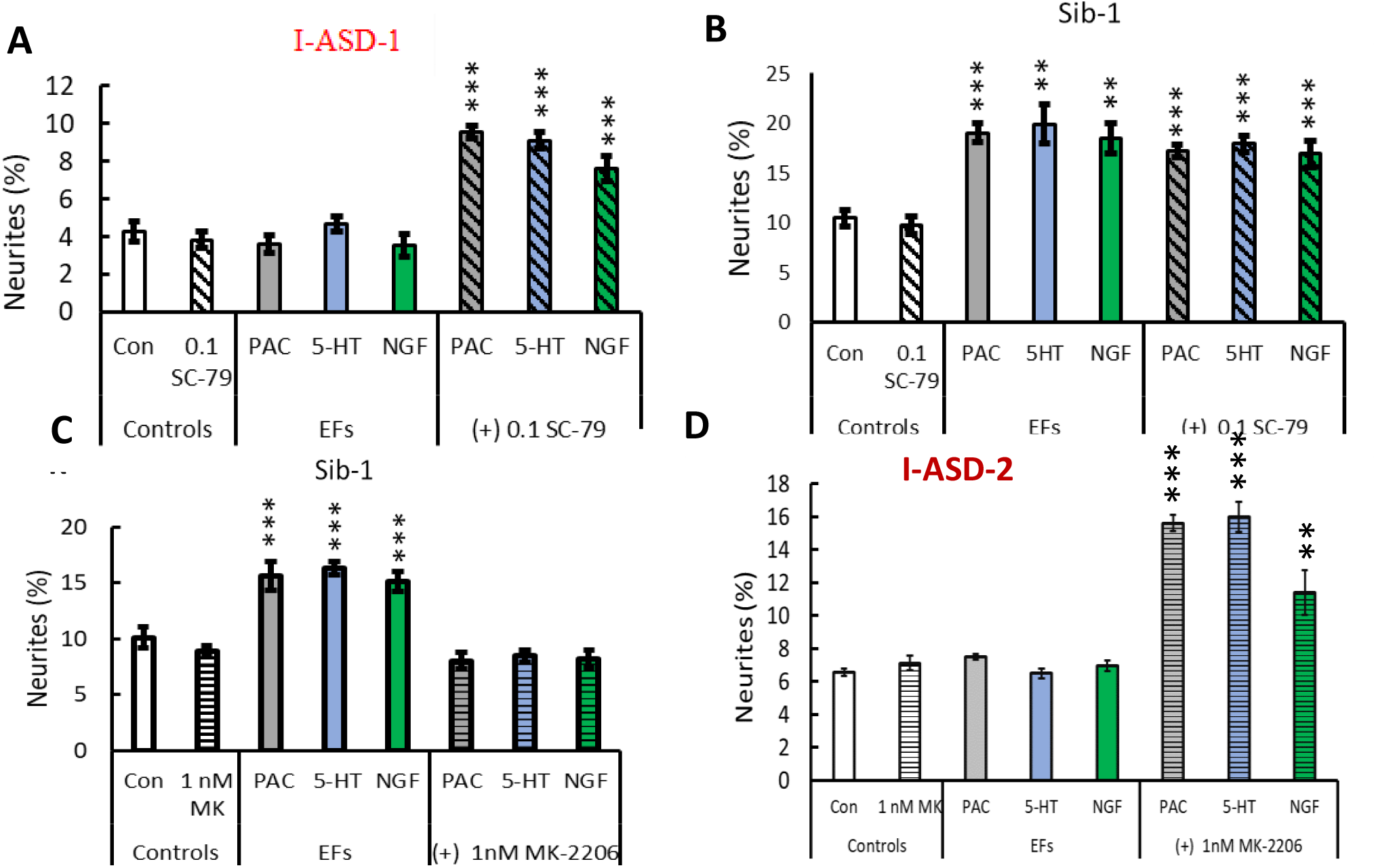
Modulation of mTOR can facilitate or abolish EF responses in I-ASD. A) Treatment of low mTOR I-ASD-1 NPCs with subthreshold dose of SC-79 (0.1 µg/mL) results in NPCs responding to EFs. B) Treatment of Sib-1 NPCs with a subthreshold dose of SC-79 does not alter EF response. C) Treatment of Sib-1 NPCs with subthreshold MK-2206 [1nM] abolishes NPC response to EFs. D) Treatment of high mTOR I-ASD-2 with subthreshold MK-2206[1 nM] establishes EF responses. P<0.001 for all comparisons, 1-way ANOVA.

Conversely, to test whether diminishing p-S6 in Sib abolishes EF responses, we took a similar “sub-threshold” approach. Addition of “sub-threshold” MK-2206 (1nM) to Sib-1 NPCs abolished their typical EF responses (Fig. 7C).

Finally, we also examined high mTOR Family-2: using a “sub-threshold” dose of inhibitor (1 nM MK-2206), I-ASD-2 NPCs were now able to respond to EFs (Fig. 7D). These “sub-threshold” studies further support altered mTOR signaling in NPCs and that manipulation of mTOR signaling can alter ASD and control NPC response to EFs during neurodevelopment.

## DISCUSSION

In this study, we utilized NPCs derived from 12 different individuals: 6 ASD cases, from two different ASD subtypes (idiopathic and 16p11.2 deletion) and 2 different control groups (Sib and NIH). Unexpectedly, we found common deficits in neurite outgrowth and cell migration in all 6 ASD NPCs. Yet, WGS revealed that I-ASD and 16pDel individuals did not share any rare variants in the 28 genes deleted in the 16p11.2 CNV and the three I-ASD individuals shared no rare functional protein coding variants that were specific to these three individuals. Despite the genetic heterogeneity, p-proteomic analyses show significant overlap between I-ASD and 16pDel with convergence onto the mTOR pathway. Subsequent western analyses validated these p-proteome differences and uncovered two distinct mTOR molecular subtypes, characterized by increased or decreased p-S6. Importantly, the mTOR differences are responsible for the shared neurodevelopmental phenotypes since manipulation of the mTOR pathway with small molecules could rescue the autism deficits and reproduce these autism phenotypes in controls. Lastly, this is one of the first studies to utilize EFs to uncover differences not apparent in control conditions which subsequently allowed for definition of two further subtypes of ASD in our dataset, EF responsive (16pDel) and EF unresponsive (I-ASD). Remarkably, mTOR manipulation also rescued EF non-responsiveness indicating the importance of this pathway as a driver of disease phenotypes. Thus, our study supports that ASD subtypes with non-overlapping genetics can have common dysregulation of mTOR signaling which is responsible for the converging common neurodevelopmental phenotypes.

### Altered Neurite Outgrowth and Migration are a Common Phenotype in All 6 ASD NPCs

While most ASD iPSC studies have investigated terminal neuronal differentiation phenotypes, our study has focused on NPCs and may be the first to find common NPC neurite and migration impairments in two different ASD subtypes. Previous iPSC-based studies of NDDs, such as Fragile X syndrome, 22q13 deletion syndrome, 16p11.2 deletion and duplication syndromes, and macrocephalic I-ASD patients have noted alterations in dendrites, axons, and neurites in post-mitotic neurons (Bhattacharyya A. et al., 2016; Castren M. et al., 2005; Doers M. E. et al., 2014; Shcheglovitov A. et al., 2013; Kathuria A. et al., 2018; Deshpande A. et al., 2017; Griesi-Oliveira K. et al., 2015; Marchetto M. C. et al., 2017). Migration defects have not been commonly noted in autism but have been reported in studies of NDDs including schizophrenia, Rett syndrome, Down Syndrome, and CNTNAP2 mutation patients (Brennand K. et al., 2015; Flaherty E. et al., 2017; Krey J. F. et al., 2013; Huo H. Q. et al., 2018; Zhang Z. N. et al., 2016).

Multiple studies indicate that NPCs are a biologically relevant cell type for ASD. Genetic studies, including extensive WES and GWAS analysis, have reproducibly identified human mid-fetal cerebral/cortical (8-24 weeks) development as a window for when and where most ASD risk genes are expressed (Satterstrom F. K. et al., 2020; Willsey A. J. et al., 2013; Parikshak N. N. et al., 2013). During this window, NPCs are formed and undergo migration and early neurite extension. Recently altered NPC development has also been identified as a consistent neurodevelopmental phenotype for numerous ASD risk genes (Garcia-Forn M. et al., 2020). Given that human iPSC-derived NPCs are similar transcriptionally and functionally to cortical NPCs, our studies provide experimental evidence of NPC developmental dysregulation in ASD (Hofrichter M. et al., 2017; Yan Y. et al., 2013a). Furthermore, by comparing two subtypes of ASD, we show, as suggested by genetics studies that this early developmental dysregulation may be a common feature amongst multiple forms of ASD.

### mTOR Signaling as a Point of Convergence in ASD

Our p-proteome, westerns and small molecule analyses identified altered mTOR signaling as a point of convergence for the shared ASD neurite and migration phenotypes. To our knowledge, we are the first to utilize unbiased p-proteomics in ASD NPCs and were able to uncover marked differences in the p-proteome for both ASD subtypes. Interestingly, despite non-overlapping genetics, significant overlap was observed (67/105) at the p-proteome level between I-ASD and 16pDel. Also, about a third of these p-proteins are common with 102 high confidence ASD risk genes or relevant gene families (Satterstrom F. K. et al., 2020), suggesting that we are identifying similar pathways as described by others. In contrast, very few proteomic changes were noted I-ASD (9/9700) and 16pDel (48/9700) when compared to unaffected individuals. This could suggest either the proteome is relatively unaltered in ASD NPCs, or the changes are quite varied between samples so no consistent significant results were observed.

The importance of mTOR signaling was further highlighted by our small molecule gain and loss of function analyses and EF studies. By correcting the P-S6 levels with mTOR agonists and antagonists, we were able to rescue the developmental defects in both I-ASD and 16pDel NPCs. Furthermore, by changing control P-S6 to ASD levels, the neurodevelopmental defects could be reproduced demonstrating that the mTOR signaling differences are mechanistically linked to the shared neurodevelopmental defects for both I-ASD and 16pDel NPCs. In addition, as one of the first studies to challenge ASD NPCs with EFs, we were able to uncover the importance of mTOR in the regulation of signaling milieu and uncover differences between I-ASD and 16pDel in their ability to respond to EFs. I-ASD NPCs failed to respond to several EFs. Yet, remarkably, in the “low mTOR” I-ASD-1 NPCs, using sub-threshold levels of the mTOR pathway activator facilitated responses to EFs while subthreshold doses of the mTOR pathway inhibitor abolished EF responses in the Sibs. Likewise in the high mTOR I-ASD-2, subthreshold levels of the inhibitor also facilitated EF response. These results suggest that mTOR homeostasis and ambient levels may be critical for facilitating several cellular processes including the ability for other signaling pathways to function effectively. As 16pDel NPCs had no EF response impairments, different ASD subtypes or different genetic backgrounds may have distinct thresholds for maintaining signaling homeostasis. Our studies point strongly to the importance of the signaling milieu and mTOR regulation in the control of neurodevelopment and ASD pathogenesis

The convergent signaling results are even more remarkable when we consider the WGS results and total proteome results. There is no rare variant overlap between the two ASD datasets, or among the three I-ASD individuals. While some I-ASD individuals have rare variants in genes that could affect mTOR signaling, these were often also shared by the Sib (ex; TSC-1 rare variants) or only individual specific and not shared amongst all three I-ASD individuals. Interestingly, assessing the genetic sequencing data from our I-ASD cohort via IPA network analysis does not reveal any enrichment in mTOR pathway molecules. Likewise, pathway analysis of published transcriptome data from 16pDel ASD patient iPSCs or lymphoblastic tissue also reveals no mTOR enrichment or dysregulation in signaling (Blumenthal I. et al., 2014). These results are also mirrored in the total proteome analyses which also do not suggest mTOR enrichment. These results suggest that even without genetic mutations or protein changes in the mTOR pathway, multiple rare and common variant genes could interact to lead to dysregulation of mTOR pathway signaling. Moreover, the presence of high and low mTOR I-ASD NPCs could conceivably be dependent upon the specific type of genetic mutations present and their effect on mTOR signaling. Consistent with this possibility, all 3 16pDel individuals who share the same CNV deletion have NPCs with high mTOR signaling. Of course, other pathways could contribute to shared neurodevelopmental defects and as indicated by dysregulation of other pathway members in p-proteome data. Ultimately, genetic analyses alone did not reveal convergence onto the mTOR pathway or alteration in the signaling milieu in our cohort. We were only able to detect our remarkably convergent mTOR deficits and alteration in signaling by utilizing p-proteomics and studying cellular signaling defects through western-blotting, further indicating that ASD could be a disorder of intracellular signaling.

Consistent with the idea that mTOR signaling dysfunction could be a common mechanism for autism, monogenic syndromic disorders, which can exhibit ASD such as Tuberous Sclerosis (TSC), PTEN-associated ASD, and Neurofibromatosis-1, have mutations that directly affect mTOR pathway components (Enriquez-Barreto L. et al., 2016; Yeung K. S. et al., 2017; Huber K. M. et al., 2015; Dasgupta B. et al., 2005; Winden K. D. et al., 2018a; Winden K. D. et al., 2019; Sharma A. et al., 2010; Rangasamy S. et al., 2016; Costales J. L. et al., 2015). Furthermore some studies of syndromic diseases where the driving mutation is not in the mTOR pathway, such as Fragile-X, Angelman, Rett Syndrome, and Phelan McDermid syndrome (22q13 deletion) also show mTOR alterations in human post mortem samples, embryonic stem cell models and hiPSC based models (Winden K. D. et al., 2018b). In 16pDel, because the MAPK3 gene is deleted, both rodent and human studies have uncovered alterations in ERK1 signaling (Pucilowska J. et al., 2018; Pucilowska J. et al., 2015; Deshpande A. et al., 2017). Importantly for 16pDel, altered mTOR has not been reported previously. On the other hand, there are no iPSC studies of I-ASD that show mTOR abnormalities. However, recent studies in peripheral white blood cells identified mTOR signaling proteins and p-proteins as being predictors of not only ASD diagnosis but also the disease severity (Onore C. et al., 2017; Rosina E. et al., 2019). Yet, it is unclear whether WBC mTOR alterations are sufficient to suggest that mTOR is contributing to neurodevelopmental phenotypes in these I-ASD individuals. Regardless, our study adds to the growing literature that mTOR signaling dysfunction is a shared pathogenic mechanism among different types of ASDs and raises the possibility that mTOR dysregulation could be observed commonly across multiple additional ASD subtypes. Thus, studying mTOR in multiple ASD subtypes could be important in deciphering the molecular etiology of ASD.

### Subtyping ASD

The heterogeneity of ASD, the subjective nature of clinical categorization, and the current inability to stratify and subtype the disorder have been postulated as reasons for the decades of unsuccessful clinical trials in ASD. Appropriately subtyping ASD would help us uncover the multiple pathways through which ASD can manifest in different individuals, allow us to take a more “personalized” approach to etiological studies, and could help us understand which therapeutics would be best suited to certain ASD subpopulations. As our study is one of the first to compare two disparate forms of ASD it has given us insight into both convergent and personalized phenotypes in ASD populations.

Despite common neurodevelopmental phenotypes, both a “low” and “high” mTOR signaling subtype is present in our ASD NPCs. Moreover, our gain and loss of function studies showed that mTOR dysfunction drives impaired neurodevelopment. Thus, characterizing ASD by mTOR levels could help us identify which groups may be candidates for mTOR activators vs inhibitors in future drug trials. Another subtype-specific difference was that I-ASD individuals failed to respond to EFs while 16pDel ASD individuals had typical increases in neurites and migration under EF stimulation. This difference in EF response did not correlate with the mTOR defects which were similar among the I-ASD-2 and 16pDel NPCs indicating that our EF studies help us glean more information than molecular studies alone. While there are no medications that target the core symptoms of ASD, we do use drugs that modulate monoamines such as serotonin (5-HT) and dopamine to manage certain behavioral symptoms in ASD. Furthermore, comorbid conditions such as depression are common in ASD and notoriously difficult to treat with our usual medications (Williams K. et al., 2010). Thus, understanding whether certain subtypes of ASD are responsive to EFs like 5-HT or even medications by using iPSC-derived cells, may help us better tailor treatments for ASD and its comorbid conditions.

### Study Limitations and Future Directions

Our study is one of the first to show the remarkable convergence of neurodevelopmental and molecular phenotypes in two distinct autism subtypes using rigorous methods such as utilization of multiple iPSC clones and neural inductions. The extensive rigor of our analyses that included a total of 29 distinct iPSC clones and 61 distinct neural inductions, lends to relatively small sample sizes (though similar to other iPSC studies) with only 6 autism probands, 3 of each subtype. Thus, it is difficult to know whether our observed phenotypes are generalizable to all individuals with 16pDel, I-ASD or other ASDs. However, it is important to note that our sample had significant clinical and genotypic heterogeneity yet, we could still uncover common phenotypes. Expanding our study design to other subtypes of ASD or other NDDs would help establish if common underpinning mechanisms could be identified (Gandal M. J. et al., 2018).

While our studies reveal important alterations in early neurodevelopment in ASD, our monolayer NPC studies do not recapitulate the 3-D nature or complex interactions occurring in the developing brain. Moreover, as hiPSC derived NPCS are most similar to fetal cortical radial glial cells, the phenotypes observed in our dishes may not directly parallel the structural or functional defects in the ASD brain. Neuropathological studies of ASD have indicated brain defects suggestive of dysregulated development. However, studies have not been conducted to determine whether, for example, a defect in migration in ASD iPSC derived NPCs is correlated with altered cortical lamination in the patient from whom the cells were derived. To understand this, MRI and postmortem analyses need to be conducted to correlate iPSC results with neuropathological alterations. Interestingly, an iPSC study of ASD patients with MRI-defined macrocephaly found that affected NPCs proliferated faster than control NPCs (Marchetto M. C. et al., 2017) suggesting correlation between structural MRI and NPC phenotypes

Our data potentially suggests that mTOR could serve as a target to employ precision medicine techniques in ASD (Sato A., 2016). However, given that NPCs that are fetal in nature, it is unclear whether targeting mTOR in a child or adult would treat disease phenotypes. Importantly, however, we know that signaling pathways such as mTOR continue to play roles in the postnatal and adult brain (Pagani M. et al., 2021). For example, mTOR is critical for the regulation of processes such as the moment-to-moment modification of dendritic spines which are essential to learning, memory, and integration of information in the brain (Ganesan H. et al., 2019; Nicolini C. et al., 2015; Tordjman S. et al., 2015; Henry F. E. et al., 2017). Thus, it is reasonable to postulate that mTOR defects, which in the prenatal period leads to altered neurites or migration could in the adult brain cause altered dendritic functioning and plasticity which could consequently lead to behavioral symptoms of ASD. As such, targeting mTOR in the adult brain could alleviate ASD symptoms. Preclinical mouse models of Rett syndrome support this concept, as treatment of adult animals with mTOR-activating IGF-1 reduced anxiety levels and increased exploratory behaviors, suggesting that targeting mTOR can have an effect outside of the fetal developmental window (Castro J. et al., 2014). On the other hand, clinical trials of IGF-1 in children with Rett syndrome and rapamycin in children with TSC have not thus far had promising effects on behavior (Khwaja O. S. et al., 2014). While early neurodevelopmental processes may no longer be occurring in the postnatal brain, studying NPCs can provide insight into the initial processes that contribute to ASD pathogenesis.

## SUPPLEMENTAL INFORMATION

**Supplementary Table S1.**
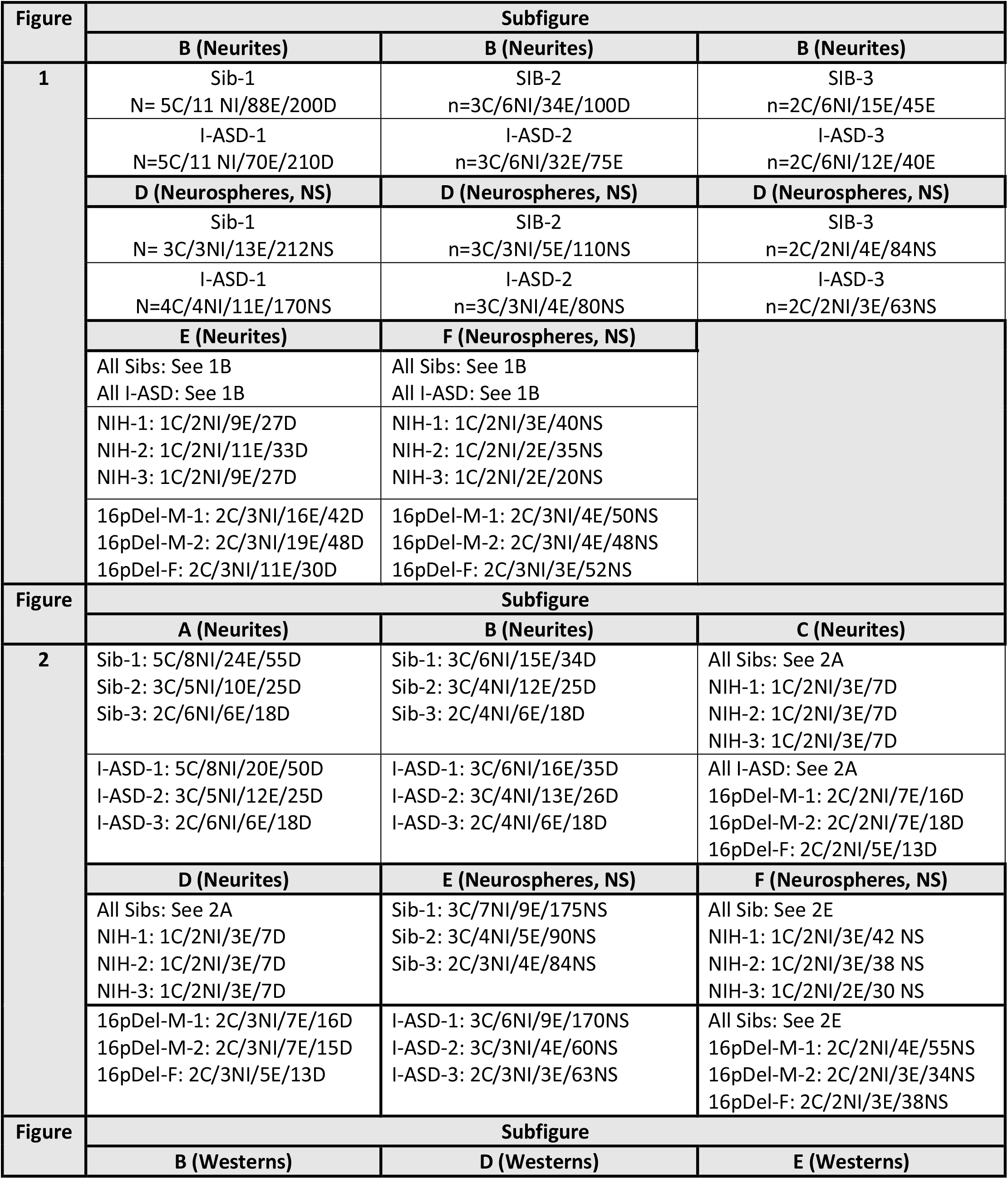
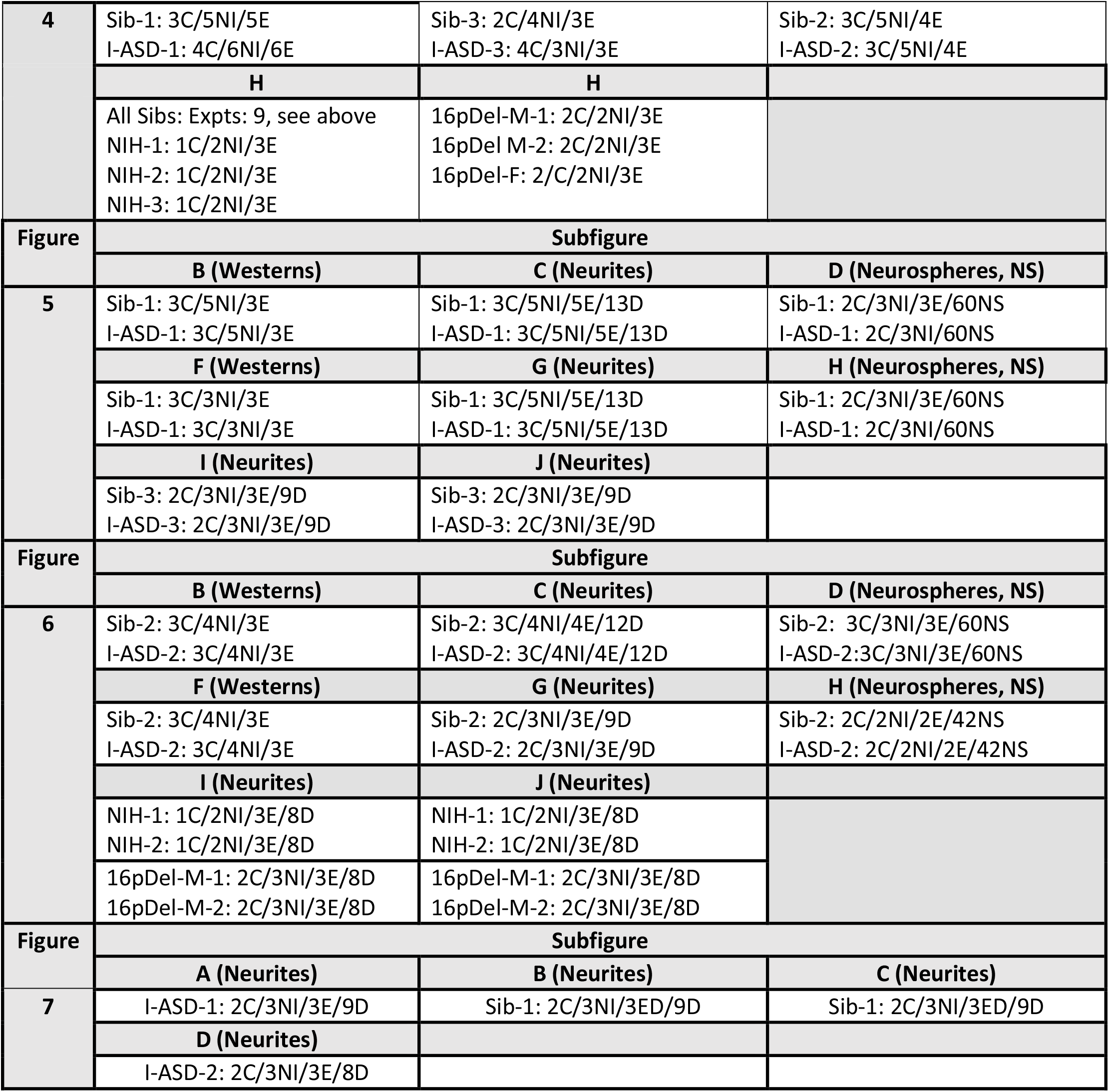
Tabulation of NPC N Values for Figures 1-7: In each cell, multiple different kinds of N-values are represented: # of clones (C)/Total # of neural inductions (NI)/ # of experiments (E) and for neurite experiments, # of dishes (D) whereas for neurospheres experiment, # of neurospheres (NS).

| Figure | Subfigure |  |  |
| --- | --- | --- | --- |
|  | B (Neurites) | B (Neurites) | B (Neurites) |
| 1 | Sib-1<br>N= 5C/11 NI/88E/200D | SIB-2<br>n=3C/6NI/34E/100D | SIB-3<br>n=2C/6NI/15E/45E |
|  | I-ASD-1<br>N=5C/11 NI/70E/210D | I-ASD-2<br>n=3C/6NI/32E/75E | I-ASD-3<br>n=2C/6NI/12E/40E |
|  | D (Neurospheres, NS) | D (Neurospheres, NS) | D (Neurospheres, NS) |
|  | Sib-1<br>N= 3C/3NI/13E/212NS | SIB-2<br>n=3C/3NI/5E/110NS | SIB-3<br>n=2C/2NI/4E/84NS |
|  | I-ASD-1<br>N=4C/4NI/11E/170NS | I-ASD-2<br>n=3C/3NI/4E/80NS | I-ASD-3<br>n=2C/2NI/3E/63NS |
|  | E (Neurites) | F (Neurospheres, NS) |  |
|  | All Sibs: See 1B<br>All I-ASD: See 1B | All Sibs: See 1B<br>All I-ASD: See 1B |  |
|  | NIH-1: 1C/2NI/9E/27D<br>NIH-2: 1C/2NI/11E/33D<br>NIH-3: 1C/2NI/9E/27D | NIH-1: 1C/2NI/3E/40NS<br>NIH-2: 1C/2NI/2E/35NS<br>NIH-3: 1C/2NI/2E/20NS |  |
|  | 16pDel-M-1: 2C/3NI/16E/42D<br>16pDel-M-2: 2C/3NI/19E/48D<br>16pDel-F: 2C/3NI/11E/30D | 16pDel-M-1: 2C/3NI/4E/50NS<br>16pDel-M-2: 2C/3NI/4E/48NS<br>16pDel-F: 2C/3NI/3E/52NS |  |
| Figure | Subfigure |  |  |
| 2 | A (Neurites) | B (Neurites) | C (Neurites) |
|  | Sib-1: 5C/8NI/24E/55D<br>Sib-2: 3C/5NI/10E/25D<br>Sib-3: 2C/6NI/6E/18D | Sib-1: 3C/6NI/15E/34D<br>Sib-2: 3C/4NI/12E/25D<br>Sib-3: 2C/4NI/6E/18D | All Sibs: See 2A<br>NIH-1: 1C/2NI/3E/7D<br>NIH-2: 1C/2NI/3E/7D<br>NIH-3: 1C/2NI/3E/7D |
|  | I-ASD-1: 5C/8NI/20E/50D<br>I-ASD-2: 3C/5NI/12E/25D<br>I-ASD-3: 2C/6NI/6E/18D | I-ASD-1: 3C/6NI/16E/35D<br>I-ASD-2: 3C/4NI/13E/26D<br>I-ASD-3: 2C/4NI/6E/18D | All I-ASD: See 2A<br>16pDel-M-1: 2C/2NI/7E/16D<br>16pDel-M-2: 2C/2NI/7E/18D<br>16pDel-F: 2C/2NI/5E/13D |
|  | D (Neurites) | E (Neurospheres, NS) | F (Neurospheres, NS) |
|  | All Sibs: See 2A<br>NIH-1: 1C/2NI/3E/7D<br>NIH-2: 1C/2NI/3E/7D<br>NIH-3: 1C/2NI/3E/7D | Sib-1: 3C/7NI/9E/175NS<br>Sib-2: 3C/4NI/5E/90NS<br>Sib-3: 2C/3NI/4E/84NS | All Sib: See 2E<br>NIH-1: 1C/2NI/3E/42 NS<br>NIH-2: 1C/2NI/3E/38 NS<br>NIH-3: 1C/2NI/2E/30 NS |
|  | 16pDel-M-1: 2C/3NI/7E/16D<br>16pDel-M-2: 2C/3NI/7E/15D<br>16pDel-F: 2C/3NI/5E/13D | I-ASD-1: 3C/6NI/9E/170NS<br>I-ASD-2: 3C/3NI/4E/60NS<br>I-ASD-3: 2C/3NI/3E/63NS | All Sibs: See 2E<br>16pDel-M-1: 2C/2NI/4E/55NS<br>16pDel-M-2: 2C/2NI/3E/34NS<br>16pDel-F: 2C/2NI/3E/38NS |
| Figure | Subfigure |  |  |
|  | B (Westerns) | D (Westerns) | E (Westerns) |
| **4** | Sib-1: 3C/5NI/5E<br>I-ASD-1: 4C/6NI/6E | Sib-3: 2C/4NI/3E<br>I-ASD-3: 4C/3NI/3E | Sib-2: 3C/5NI/4E<br>I-ASD-2: 3C/5NI/4E |
|  | **H** | **H** |  |
|  | All Sibs: Expts: 9, see above<br>NIH-1: 1C/2NI/3E<br>NIH-2: 1C/2NI/3E<br>NIH-3: 1C/2NI/3E | 16pDel-M-1: 2C/2NI/3E<br>16pDel M-2: 2C/2NI/3E<br>16pDel-F: 2/C/2NI/3E |  |
| **Figure** | **Subfigure** |  |  |
|  | **B (Westerns)** | **C (Neurites)** | **D (Neurospheres, NS)** |
| **5** | Sib-1: 3C/5NI/3E<br>I-ASD-1: 3C/5NI/3E | Sib-1: 3C/5NI/5E/13D<br>I-ASD-1: 3C/5NI/5E/13D | Sib-1: 2C/3NI/3E/60NS<br>I-ASD-1: 2C/3NI/60NS |
|  | **F (Westerns)** | **G (Neurites)** | **H (Neurospheres, NS)** |
|  | Sib-1: 3C/3NI/3E<br>I-ASD-1: 3C/3NI/3E | Sib-1: 3C/5NI/5E/13D<br>I-ASD-1: 3C/5NI/5E/13D | Sib-1: 2C/3NI/3E/60NS<br>I-ASD-1: 2C/3NI/60NS |
|  | **I (Neurites)** | **J (Neurites)** |  |
|  | Sib-3: 2C/3NI/3E/9D<br>I-ASD-3: 2C/3NI/3E/9D | Sib-3: 2C/3NI/3E/9D<br>I-ASD-3: 2C/3NI/3E/9D |  |
| **Figure** | **Subfigure** |  |  |
|  | **B (Westerns)** | **C (Neurites)** | **D (Neurospheres, NS)** |
| **6** | Sib-2: 3C/4NI/3E<br>I-ASD-2: 3C/4NI/3E | Sib-2: 3C/4NI/4E/12D<br>I-ASD-2: 3C/4NI/4E/12D | Sib-2: 3C/3NI/3E/60NS<br>I-ASD-2: 3C/3NI/3E/60NS |
|  | **F (Westerns)** | **G (Neurites)** | **H (Neurospheres, NS)** |
|  | Sib-2: 3C/4NI/3E<br>I-ASD-2: 3C/4NI/3E | Sib-2: 2C/3NI/3E/9D<br>I-ASD-2: 2C/3NI/3E/9D | Sib-2: 2C/2NI/2E/42NS<br>I-ASD-2: 2C/2NI/2E/42NS |
|  | **I (Neurites)** | **J (Neurites)** |  |
|  | NIH-1: 1C/2NI/3E/8D<br>NIH-2: 1C/2NI/3E/8D | NIH-1: 1C/2NI/3E/8D<br>NIH-2: 1C/2NI/3E/8D |  |
|  | 16pDel-M-1: 2C/3NI/3E/8D<br>16pDel-M-2: 2C/3NI/3E/8D | 16pDel-M-1: 2C/3NI/3E/8D<br>16pDel-M-2: 2C/3NI/3E/8D |  |
| **Figure** | **Subfigure** |  |  |
|  | **A (Neurites)** | **B (Neurites)** | **C (Neurites)** |
| **7** | I-ASD-1: 2C/3NI/3E/9D | Sib-1: 2C/3NI/3ED/9D | Sib-1: 2C/3NI/3ED/9D |
|  | **D (Neurites)** |  |  |
|  | I-ASD-2: 2C/3NI/3E/8D |  |  |

**Supplementary Table S2.** Alternative Allele Genotypes in the chr16.p11.2 Deletion Region in 3 I-ASD Families: The count of the heterozygous and homozygous alternative allele genotypes and their ratio in the chr16.p11.2 deletion region (chr16: 28,500,001-35,300,000). In the event of a chr16.p11.2 deletion, we expect no heterozygous genotypes in the region. The large number of heterozygous genotypes across all individuals in this region indicates that the deletion does not appear in any of the individuals in the 3 I-ASD families.

| **Sequenced Individual** | **# of heterozygous alternative genotypes** | **# of homozygous alternative genotypes** | **% of alternative heterozygous genotypes** |
| --- | --- | --- | --- |
| <u>Family-1 Male Parent</u><br>SL128062 | 15889 | 20041 | 44.2% |
| <u>Family-1 Female Parent</u><br>SL128065 | 12890 | 23040 | 35.9% |
| <u>Family-1 LLI</u><br>07C69064 | 11875 | 24055 | 33.1% |
| <u>I-ASD-1</u><br>2743-SL-0029 | 9224 | 26706 | 25.7% |
| <u>Sib-1</u><br>07C69065 | 15305 | 20625 | 42.6% |
| <u>Family-2 Male Parent</u><br>SL128102 | 17859 | 18071 | 49.7% |
| <u>Family-2 Female Parent</u><br>SL128091 | 14600 | 21330 | 40.6% |
| <u>I-ASD-2</u><br>07C65371 | 15688 | 20242 | 43.7% |
| <u>Sib-2</u><br>07C66696 | 15536 | 20394 | 43.2% |
| <u>Family-3 Male Parent</u><br>SL127748 | 16822 | 19108 | 46.8% |
| <u>Family-3 Female Parent</u><br>SL127734 | 14769 | 21161 | 41.1% |
| <u>Family-3 LLI</u><br>SL127742 | 16785 | 19145 | 46.7% |
| <u>Sib-F</u><br>05C40413 | 14687 | 21243 | 40.9% |
| <u>Sib-3</u><br>05C40411 | 15140 | 20790 | 42.1% |
| <u>I-ASD-3</u><br>2743-SL-0028 | 11341 | 24589 | 31.6% |

**Figure S1.**
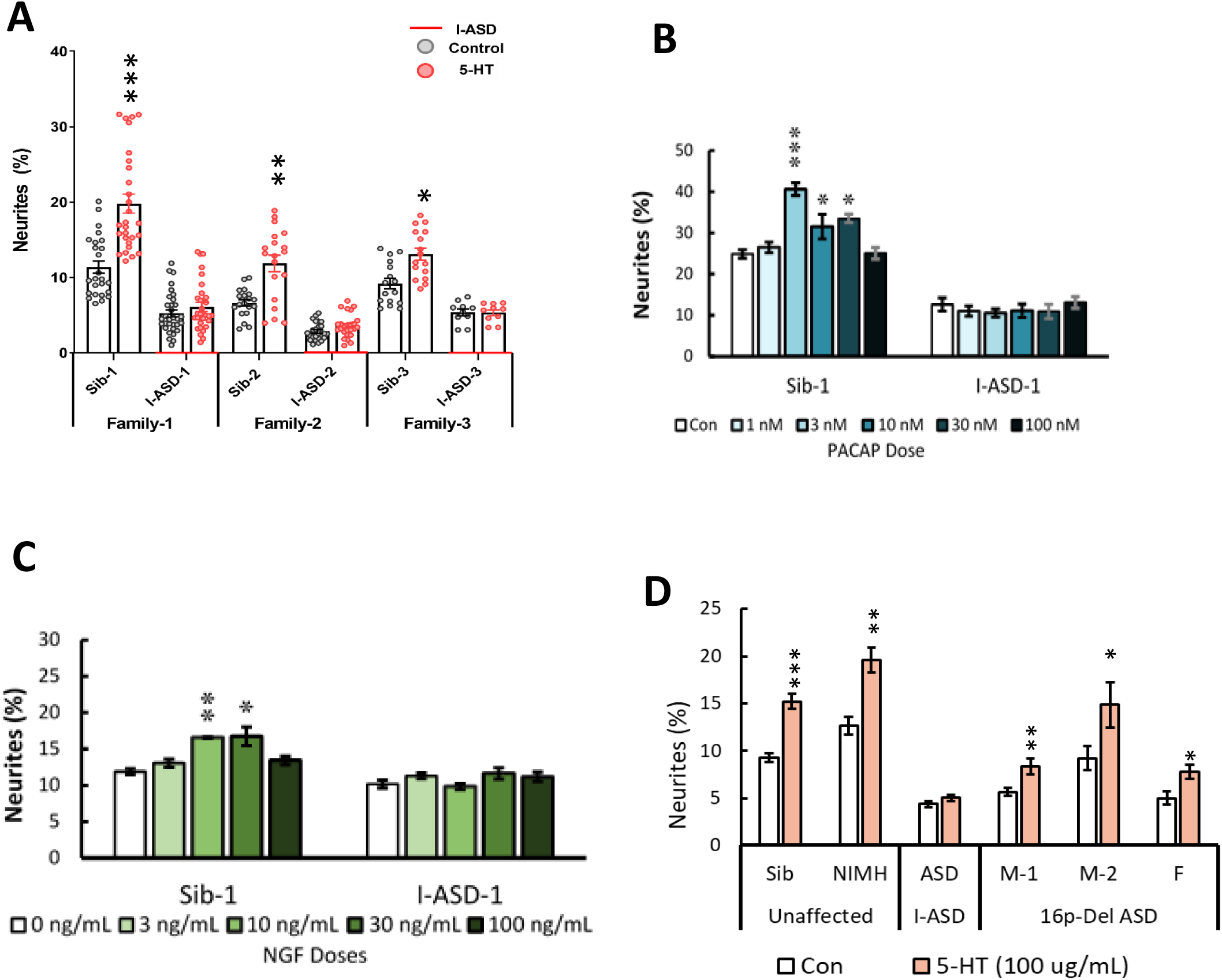
Extended PACAP, 5HT, and NGF EF studies. A) 5-HT increases neurite outgrowth in I-ASD Sib NPCs but has no effect on I-ASD affected NPCs (Sib vs ASD comparisons-Student’s T-test, p<0.001). B) Dose response experiments-PACAP: Sib is responsive to PACAP at numerous concentrations whereas I-ASD has no response at any concentration (p<0.001 at all doses Sib vs I-ASD, 1 way ANOVA). C) Dose response experiments-NGF: Sib is responsive to NGF at numerous concentrations whereas I-ASD has no response at any concentration (p<0.001 at all doses Sib vs I-ASD, 1 way ANOVA). D) 5-HT stimulates neurite outgrowth in I-ASD Sib, NIH, and all 3 16pDel NPCs but not in I-ASD affected NPCs (2-way ANOVA).

**Figure S2.**
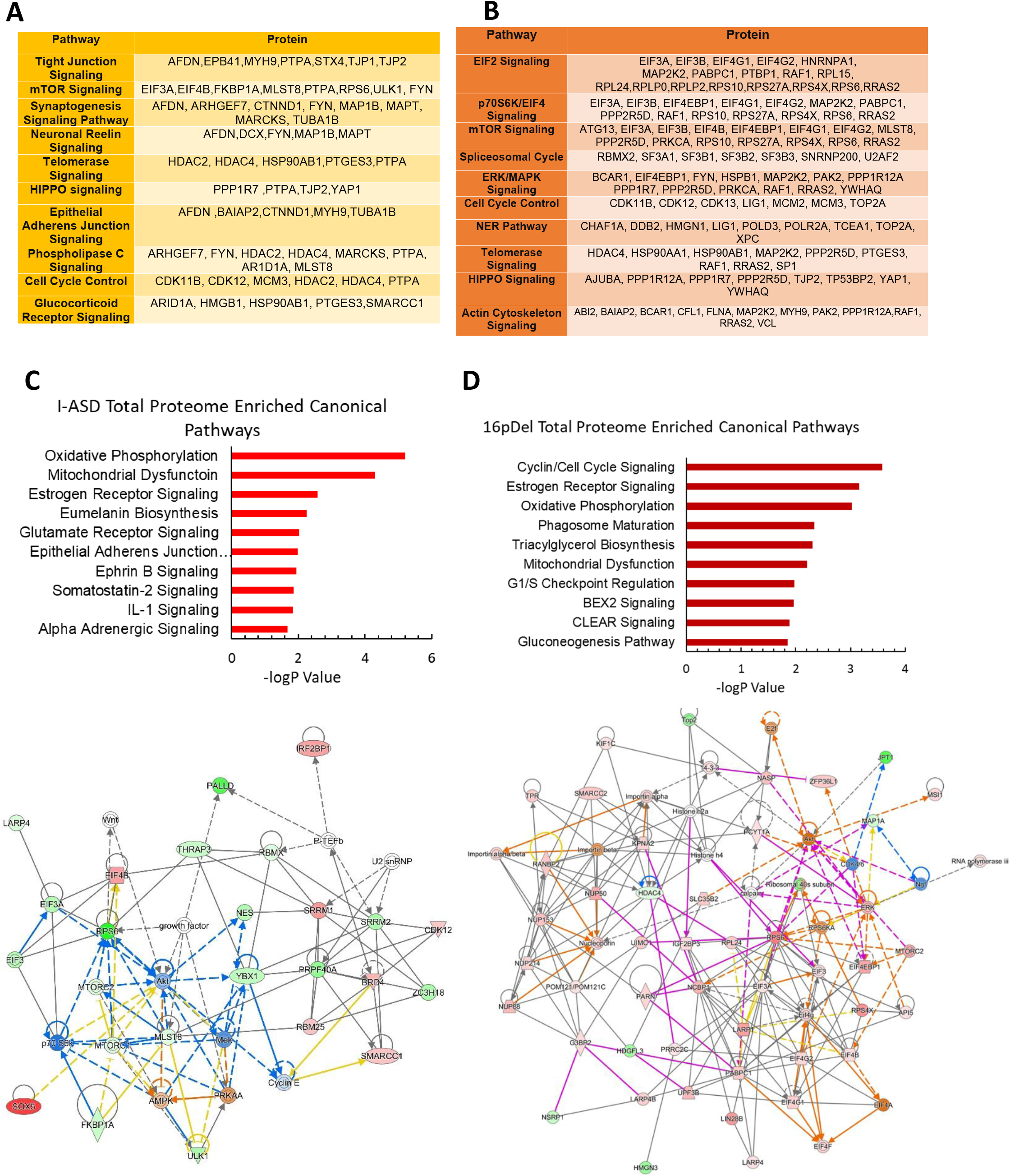
Further Analyses of the P-Proteome. A) Table showing the protein members in each enriched canonical pathway in I-ASD B) Table showing the protein members in each enriched canonical pathway in 16pDel ASD C) IPA canonical pathway analysis of total proteome data from I-ASD showing top 10 enriched pathways. Total proteome does not show enrichment of mTOR. D) IPA canonical pathway analysis of total proteome data from 16pDel showing top 10 enriched pathways. Total proteome does not show enrichment of mTOR. E) Network analysis of I-ASD p-proteome showing one of the most enriched networks. mTOR and associated pathway members (AKT, RPS6, EIFs) are highly enriched particularly looking at the left side of the image. F) Network analysis of 16pDel p-proteome showing one of the most enriched networks. mTOR pathway members (RPS6, EIF, PS60K, AKT) are highly enriched, particularly looking at the right side of the network.

**Figure S3.**
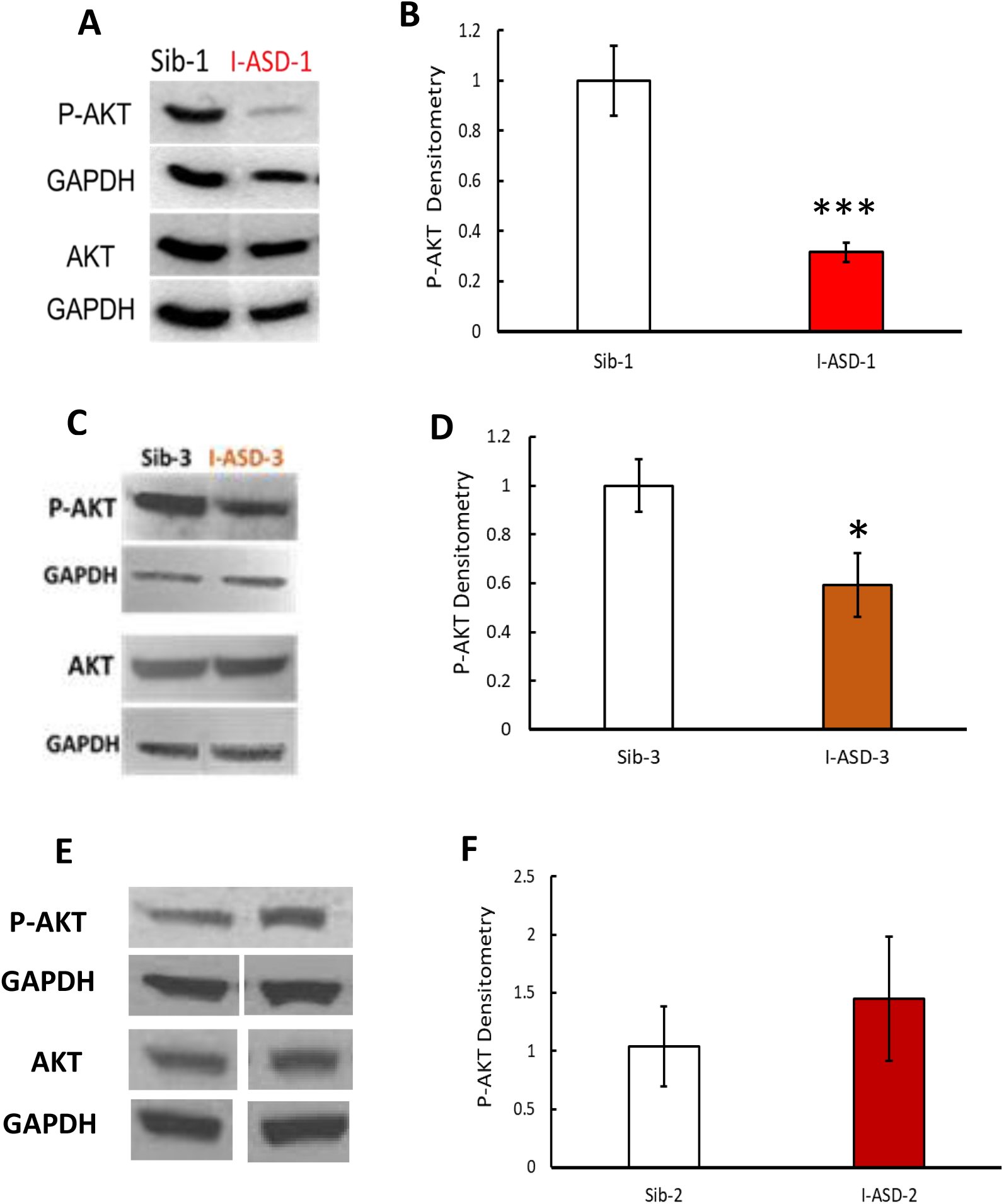
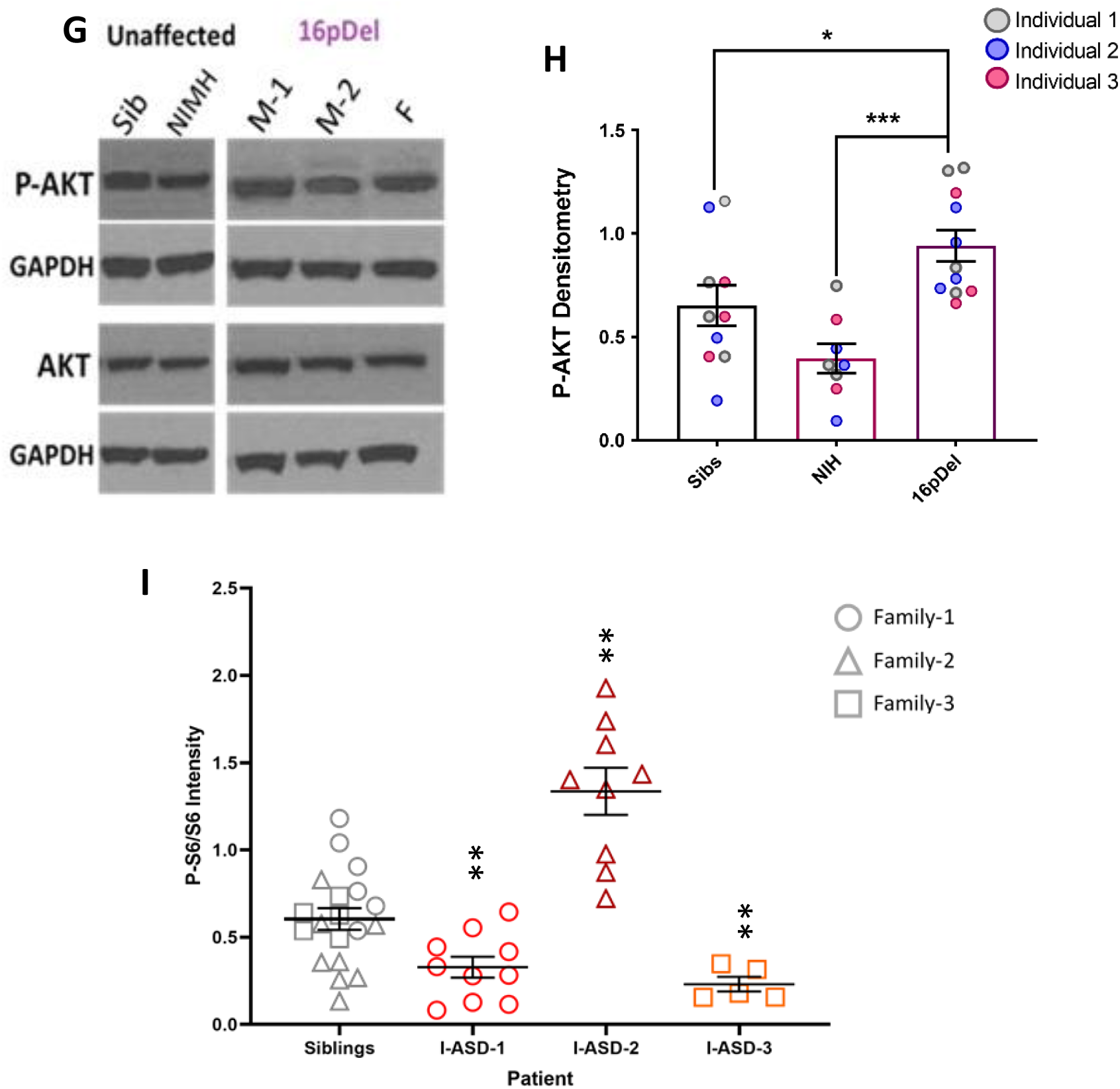
P-AKT in I-ASD and 16pdel NPCs. All images show P-AKT and AKT western blots with matched GAPDH loading control. Graphs show densitometry quantifications of normalized P-AKT (P-AKT/GAPDH) divided by normalized AKT (AKT/GAPDH). Student’s T-test was used for all I-ASD comparisons. A) I-ASD-1 representative western blot showing reduced P-AKT but similar AKT and GAPDH in I-ASD-1 compared to Sib-1 B) Graph: reduced P-AKT/AKT in I-ASD-1 vs. Sib-1 (p<0.0001) C) I-ASD-3 Representative western blot showing reduced P-AKT, but similar AKT and GAPDH in I-ASD-3 vs. Sib-3 (p<0.0001). D) Graph: Reduced P-AKT/AKT levels in I-ASD-3 compared to Sib-3. E) Representative western blot showing slightly increased P-AKT but similar AKT and GAPDH in I-ASD-2 compared to Sib-2. F) Graph: levels in I-ASD-2 demonstrate a significant trend towards an increase in p-AKT compared to Sib-2 (p<0.084). G) Western blot showing increased P-AKT in all 16pDel NPCs compared to both NIH and Sib with approximately equal total AKT and GAPDH. H) Graph: Increased P-AKT/AKT in all 16pdel NPCs compared to Sib and NIH NPCs (1-way ANOVA, p<0.001). I) Comparison of P-S6 levels between all I-ASD Sib controls (all sibs) and each I-ASD affected individual demonstrates that I-ASD-1 and 3 have reduced P-S6 while I-ASD-2 has elevated P-S6, indicating that phenotypes are relevant even when compared to all I-ASD Sib controls non-Sibs.

**Figur S4.**
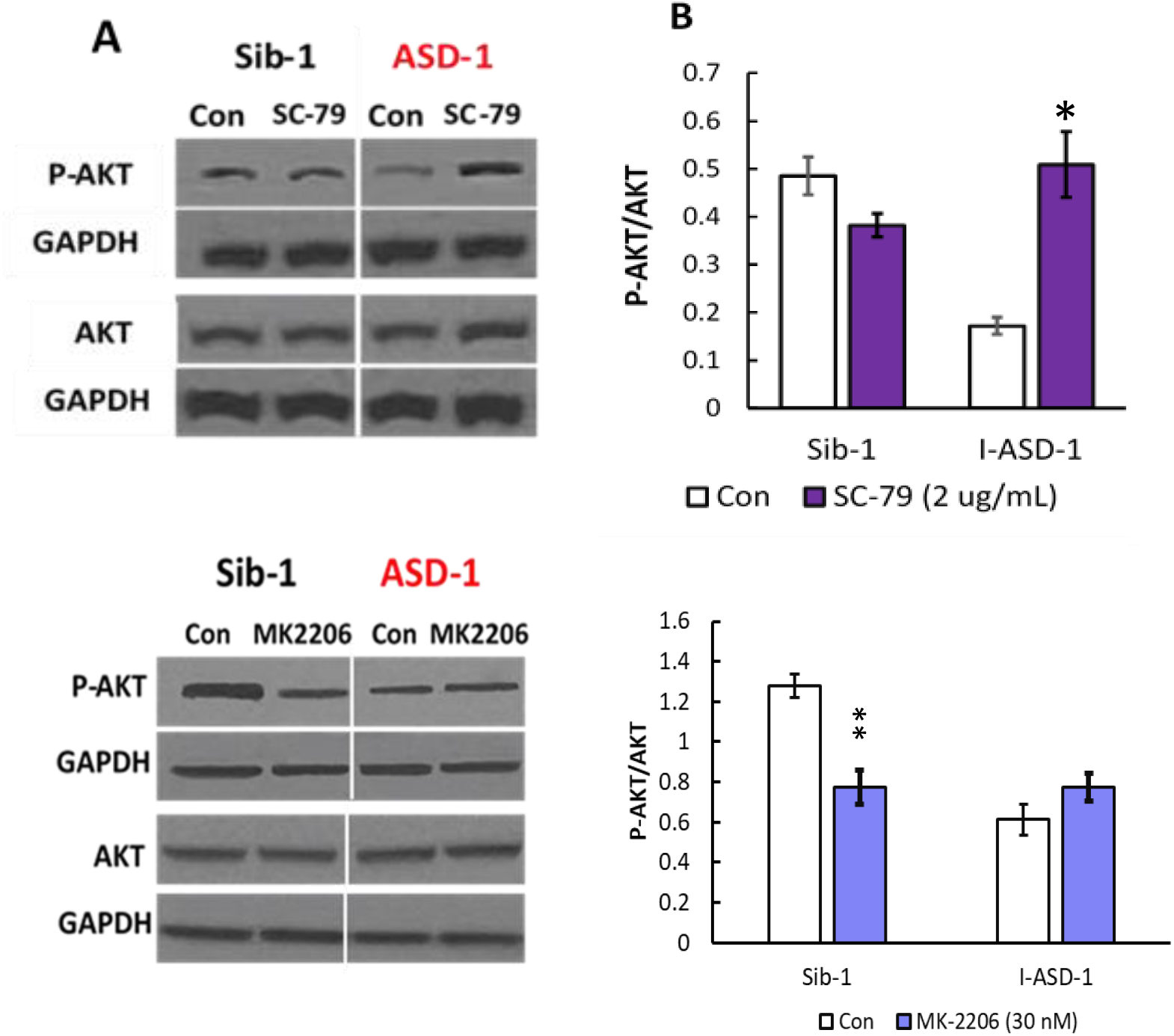
P-AKT changes with SC-79 and MK-2206 in Family-1. All images show P-AKT and AKT western blots with matched GAPDH loading control. Graphs show densitometry quantifications of normalized P-AKT (P-AKT/GAPDH) divided by normalized AKT (AKT/GAPDH). Student’s T-test was used for all comparisons. A) Representative blot showing that SC-79 does increased P-AKT but not AKT levels in I-ASD-1 with no changes to Sib-1 P-AKT or AKT B) Graph: Quantification of P-AKT/AKT increase in I-ASD-1 with SC-79 (p<0.01) with no changes in Sib P-AKT/AKT (p=0.21) C) Representative blot showing that MK-2206 (30 nM) does reduce P-AKT but not AKT levels in Sib-1 with no changes in I-ASD-1 P-AKT or AKT levels. D) Graph: Quantification showing that MK-2206 reduces P-AKT/AKT levels in Sib-1 (P<0.001) without changing P-AKT/AKT in I-ASD-1 (p=0.63).

## ACKNOWLEDGEMENTS

Our sincerest thanks to the NJLAGs families whose interest in supporting autism research allowed for the findings in our studies and Drs. Linda Brzustowicz and Judy Flax for their collaboration and support. We also give our sincerest thanks to the families participating in the Simons Searchlight Sites as well as the Simons Searchlight Consortium. We appreciate obtaining access to the phenotypic data and the cellular biospecimens on the SFARI base as well. We thank RUCDR Infinite Biologics for providing Simons Searchlight 16p11.2 Deletion and NIH Regenerative Medicine Program iPSCs as well as for their support and technical guidance throughout the study. Our thanks to the Center for Integrative Proteomics Research and the Mass Spectrometry Facility of Rutgers Robert Wood Johnson Medical School for their support and technical guidance for the proteomic analyses. We would like to thank the following individuals for their contributions to this manuscript: Courtney McDermott, Maya Hale, Katelyn Jo, Angelique Sarrosquy, and Meghan Eller.

We would like to acknowledge the following funding sources as well. This work was supported by the New Jersey Governor’s Council for Medical Research and Treatment of Autism (CAUT13APS010, CAUT14APL031, CAUT15APL041, CAUT19APL014) and the Nancy Lurie Marks Family Foundation for E.D.-B. and J.H.M.; the NJ Health Foundation (PC 63-19) for J.H.M.; the Mindworks Charitable Lead Trust and Jewish Community Foundation of Greater MetroWest for E.D.-B.; an Autism Science Foundation Undergraduate summer research grant for C.P. and E.D.-B. and Rutgers School of Graduate Studies for S.P. and E.D.-B. J.X. was supported by the New Jersey Governor’s Council for Medical Research and Treatment of Autism grant CAUT19APL028.”

## AUTHOR CONTRIBUTIONS

Conceptualization: E.D.B., J.H.M., S.P., J.X.

Methodology: E.D.B., J.H.M., S.P., P.M.

Formal Analysis: S.P., M.M., R.A.

Investigation: S.P., B.D., C.P., M.M., R.A., R.J.C., M.S.T., X.Z., P.M.

Writing: S.P., E.D.B., J.H.M.

Visualization: S.P, R.J.C., C.P., B.D., M.S.T., M.M.

NPC Line Generation: S.P., M.S.T., R.J.C., M.M., P.M.

Funding Acquisition: E.D.B., J.H.M., J.X.

Supervision: E.D.B, J.H.M, J.X.

